# Viral manipulation of mechanoresponsive signaling disassembles processing bodies

**DOI:** 10.1101/2020.05.15.091876

**Authors:** Elizabeth L. Castle, Carolyn-Ann Robinson, Pauline Douglas, Kristina D. Rinker, Jennifer A. Corcoran

## Abstract

Processing bodies (PBs) are ribonucleoprotein granules that suppress cytokine mRNA translation that are targeted for disassembly by many viruses. Kaposi’s sarcoma-associated herpesvirus is the etiological agent of the inflammatory endothelial cancer, Kaposi’s sarcoma, and a PB-regulating virus. The virus encodes Kaposin B (KapB), which induces actin stress fibres (SFs) and cell spindling as well as PB disassembly. We now show that KapB-mediated PB disassembly requires actin rearrangements, RhoA effectors and the mechanoresponsive transcription activator, YAP. Moreover, ectopic expression of active YAP or exposure of ECs to mechanical forces caused PB disassembly in the absence of KapB and mechanoresponsive PB disassembly also required YAP. Using the viral protein KapB, we identified a new consequence of the exposure of cells to mechanical forces that alter actin dynamics and activate YAP, namely the disassembly of PBs.

**Importance:** For the first time, we demonstrate that processing bodies (PBs), cytoplasmic sites of RNA decay, are regulated by mechanical signaling events that alter actin dynamics and that this requires the mechanoresponsive transcription factor, YAP. Using the overexpression of a viral protein called KapB, known previously to mediate PB disassembly, we show that actin stress fibers (SFs) and the mechanoresponsive transcription factor, YAP, are required for PB loss. We also show that other established mechanical signals (shear stress or stiff extracellular matrix) that lead to the formation of SFs and activate YAP also cause PB disassembly. This is important because it means that KapB activates, from the inside out, a pathway that links cell shape to post-transcriptional gene regulation via cytoplasmic PBs.

## Introduction

Cells are exposed to a variety of environments and they respond to changes in external force by adjusting internal tension. These mechanical cues can be transmitted to the cell through changes to extracellular contact nodes (focal adhesions) and contractile actomyosin structures to maintain tension homeostasis (Friedland, Lee, and Boettiger 2009; Kong et al. 2009; del Rio et al. 2009; Grashoff et al. 2010; reviewed in Finch-Edmondson and Sudol 2016). Actin stress fibres (SFs) are cytoskeletal structures composed of thick actin bundles, often associated with focal adhesions (Vallenius 2013), that are force-responsive, maintaining cytoskeletal integrity in changing mechanical environments (Burridge and Guilluy 2016). SF formation is coordinated by the GTPase, RhoA; it activates the formin, mammalian diaphanous protein-1 (mDia1) to promote actin filament growth and Rho-associated coiled-coil kinase (ROCK) to promote actomyosin contractility through non-muscle myosin II (Watanabe et al. 1997; Amano et al. 1997; Kimura et al. 1996). These RhoA-effectors act together to promote the formation of contractile and stable actin filaments in response to mechanical and chemical stimuli (Watanabe et al. 1999).

External forces elicit a cascade of signals using actin as force transducers to alter gene expression. The activity of the mechanoresponsive transcriptional coactivator, Yes-associated protein (YAP), is controlled by cell shape and cytoskeletal structure (Dupont et al. 2011; Wada et al. 2011; Halder, Dupont, and Piccolo 2012; Yu et al. 2012). YAP is nuclear and active in response to low cell-cell contact (Zhao et al. 2007), high stiffness of the extracellular matrix (ECM) (Dupont et al. 2011), in shear stress due to fluid flow (K.-C. Wang et al. 2016; Nakajima et al. 2017; Lai and Stainier 2017; H. J. Lee et al. 2017; Huang et al. 2016), or after G-protein coupled receptor (GPCR) activation (Yu et al. 2012). Most of these signals induce the activity of RhoA and promote the formation of SFs, and studies have implicated actin cytoskeletal structures as requisite intermediates for YAP activation (Noria et al. 2004; Lee and Kumar 2016).

Nuclear YAP associates with its coactivators to mediate transcription of genes involved in cell proliferation, differentiation, survival and migration (Halder, Dupont, and Piccolo 2012). Consistent with this, nuclear YAP is often pro-tumourigenic and drives progression of many oncogenic traits in a variety of cancers. These include the induction of cell stemness (Panciera et al. 2016), altered metabolism (C. Yang et al. 2018), cancer cell invasion/vascular remodeling (Calvo et al. 2013; Liu et al. 2018; Kimura et al. 2020), and altered growth and proliferation (Kapoor et al. 2014; Zanconato et al. 2015; Jang et al. 2017).

Kaposi’s sarcoma (KS) is an endothelial cell (EC) cancer that is caused by the virus, Kaposi’s sarcoma-associated herpesvirus (KSHV) (Chang et al. 1994; Russo et al. 1996; Zhong et al. 1996; Ganem 1997). KSHV establishes persistent, life-long infection of its human host, and displays two types of infection, latent and lytic. In KS, the majority of the tumour ECs are latently infected while lytic replication is rare; in part, because these cells die as a result of viral replication (Boshoff et al. 1995; Staskus et al. 1997; Umbach et al. 2010; Speck and Ganem 2010). KS tumour ECs elongate into a ‘spindled’ morphology and expel proinflammatory cytokines and angiogenic factors, which promote tumour progression and create a highly inflammatory tumour microenvironment. Many features of *in vivo* KS are recapitulated by *in vitro* latent infection of primary ECs, or ectopic expression of individual KSHV latent genes, including spindling and pro-inflammatory characteristics (Ensoli 1998; Ciufo et al. 2001; Naranatt et al. 2003; Grossmann et al. 2006; Ojala and Schulz 2014; DiMaio et al. 2011). Both spindling and proinflammatory gene expression can induced by two different KSHV latent genes, vFLIP (Grossmann et al. 2006) and Kaposin B (KapB; (Corcoran, Johnston, and McCormick 2015)), suggesting a linkage between these two phenotypes. Furthermore, during their short lifetime, lytically infected cells also secrete large quantities of pro-inflammatory and angiogenic molecules. A contributor to this secretory phenotype is the constitutively active viral G protein-coupled receptor (vGPCR), a lytic viral protein that also promotes cell elongation, RhoA activation and cytoskeletal remodeling (Shepard et al. 2001; Montaner et al. 2006; Corcoran et al. 2012). vGPCR also activates YAP during KSHV lytic infection (Liu et al. 2015); however, no information exists to demonstrate YAP involvement in KSHV latency.

One way that KSHV latency promotes the pro-inflammatory and pro-tumourigenic KS microenvironment is via KapB-mediated disassembly of cytoplasmic ribonucleoprotein granules called processing bodies (PBs) (Corcoran, Johnston, and McCormick 2015). PBs are involved in many RNA regulatory processes such as RNA silencing, nonsense-mediated decay and mRNA decay and translational suppression of mRNA (Eulalio, Behm-Ansmant, and Izaurralde 2007, Horvathova et al. 2017, Tutucci et al. 2018, Wilbertz et al. 2019, Hubstenberger et al. 2017). We and others have shown that PBs are the major site for the translational suppression or constitutive decay of human mRNAs that code for potent regulatory molecules such as proinflammatory cytokines (Franks and Lykke-Andersen 2007; Corcoran, Johnston, and McCormick 2015; Vindry et al. 2017; Blanco et al. 2014). There are ∼4500 of these transcripts, all of which bear destabilizing AU-rich elements (AREs) in their 3’-untranslated regions (3’-UTRs) (Shaw and Kamen 1986; Shyu, Greenberg, and Belasco 1989; Chen and Shyu 1995; Winzen et al. 1999; Stoecklin, Mayo, and Anderson 2006; Franks and Lykke-Andersen 2007; Bakheet, Williams, and Khabar 2006; Bakheet, Hitti, and Khabar 2017). PB abundance and composition is extremely dynamic and responds to cellular stress (Sheth 2003; Kedersha and Anderson 2007; Aizer et al. 2008; Takahashi et al. 2011). Specifically, activation of the stress-responsive p38/MK2 MAP kinase pathway by KapB elicits PB disassembly and prevents constitutive ARE-mRNA turnover or translation suppression (Winzen et al. 1999; Docena et al. 2010; Corcoran et al. 2012; Corcoran and McCormick 2015; Corcoran, Johnston, and McCormick 2015). This is an important yet underappreciated regulatory mechanism that fine tunes the production of potent proinflammatory cytokines and angiogenic factors in KS.

Though PBs are dynamic and stress-responsive, the precise signaling events that lead to PB assembly or disassembly are not well understood. We showed previously that KapB binds and activates MK2, which then phosphorylates hsp27, complexes with p115RhoGEF, and activates RhoA to elicit PB disassembly (Corcoran, Johnston, and McCormick 2015; Garcia et al. 2009; McCormick and Ganem 2005). While it is well-established that RhoA coordinates SF formation (Ridley and Hall 1992; Watanabe et al. 1999; Schmitz et al. 2000; Hotulainen and Lappalainen 2006), the precise mechanism of how RhoA promotes PB disassembly is not appreciated (Corcoran, Johnston, and McCormick 2015; Takahashi et al. 2011). In an effort to better understand the regulation of PB disassembly by KapB and RhoA, we began by targeting downstream RhoA effectors reported to promote SF formation to determine if the proteins known to mediate cytoskeletal remodeling were also necessary for PB disassembly. We reasoned that at some point we would be able to uncouple the signaling events that led to SFs from those that led to PB disassembly. We were not. We now present data that conclusively shows KapB-mediated PB disassembly is dependent not only on RhoA, but on cytoskeletal structures, actomyosin contractility and the presence of the mechanoresponsive transcription transactivator, YAP. We also present the first evidence of the involvement of YAP in the tumourigenic phenotypes induced by a KSHV latent gene, KapB. We also extend these studies beyond their impact on viral tumourigenesis, by determining the mechanical regulation of PB dynamics in the absence of KapB expression, and show that induced cell contractility, cytoskeletal structures and active YAP cause PB disassembly. Using a viral protein from an oncogenic virus, we have discovered a mechanoresponsive signaling pathway that transduces signals from cell shape and cytoskeletal structures to YAP to control PBs, post-transcriptional regulators of cellular gene expression.

## Results

### RhoA effectors controlling SF formation are required for PB disassembly

We previously showed that KapB-mediated PB disassembly required RhoA (Corcoran, Johnston, and McCormick 2015). In this work, we investigated whether downstream RhoA effectors known to control SF formation also control PB disassembly. Mammalian diaphanous protein 1 (mDia1) and Rho-associated coiled-coil kinase (ROCK) are considered the main coordinators of RhoA-mediated SF formation (Watanabe et al. 1999; Tojkander, Gateva, and Lappalainen 2012). mDia1 is a formin that promotes actin filament polymerization (Watanabe et al. 1999). To examine whether mDia1 was required for KapB-mediated PB disassembly, we designed short hairpin RNAs (shRNAs) to silence mDia1 mRNA. KapB- and vector-expressing human umbilical vein endothelial cells (HUVECs) were transduced with mDia1-targeting shRNAs and selected. Silencing efficacy was confirmed with immunoblotting (Fig 1A). PB analysis was performed using CellProfiler to quantify immunofluorescence images stained for the hallmark PB-resident protein, Enhancer of mRNA decapping 4 (EDC4), as described in detail in the methods (J. H. Yu et al. 2005; Kedersha et al. 2008). Knockdown of mDia1 increased PBs in KapB-expressing cells (Fig 1B, D). mDia1-sh1 showed a greater increase in PBs in comparison to mDia1-sh2 (Fig 1B), and also increased PBs in vector control cells, likely because mDia1-sh1 reduced protein expression by 90% whereas mDia1-sh2 reduced it by 40- 50% (Fig 1A). To separate how silencing of mDia1 impacted baseline turnover of PBs from how silencing impacts KapB-mediated PB disassembly, we calculated the ratio of PBs per cell in KapB-expressing cells and normalized to PBs per cell in the control. This is important because this calculation shows whether KapB is still able to disassemble PBs, relative to vector, in the context of mDia1 silencing. If the ratio is ≥1 after sh-mDia1 treatment, it indicates that KapB is no longer able to disassemble PBs in comparison to the vector control, and that mDia1 contributes directly to KapB-mediated PB disassembly. Conversely, if the ratio is ∼ 0.4 to 0.6, it indicates that KapB can still disassemble PBs even in the context of sh-mDia1 treatment. In this case, we determined that silencing using both mDia1-sh1 and mDia1-sh2 restored the PB ratio in KapB:Vector cells to ∼1, indicating that the ability of KapB to disassemble PBs is lost after mDia1 silencing and that this is a KapB-specific effect (Fig 1C). We note that this ratio will be reported in subsequent figures for every RNA silencing or drug treatment applied to test KapB-mediated PB disassembly. We also observed that mDia1 silencing did not eliminate SF formation (Fig 1D) but, instead, increased elongated cells with visible actin SFs across the cell in both vector and KapB conditions. The visible actin structures may represent different SF subtypes or actin bundles that compensate for the loss of mDia1 (Hotulainen and Lappalainen 2006).

**Figure 1:**
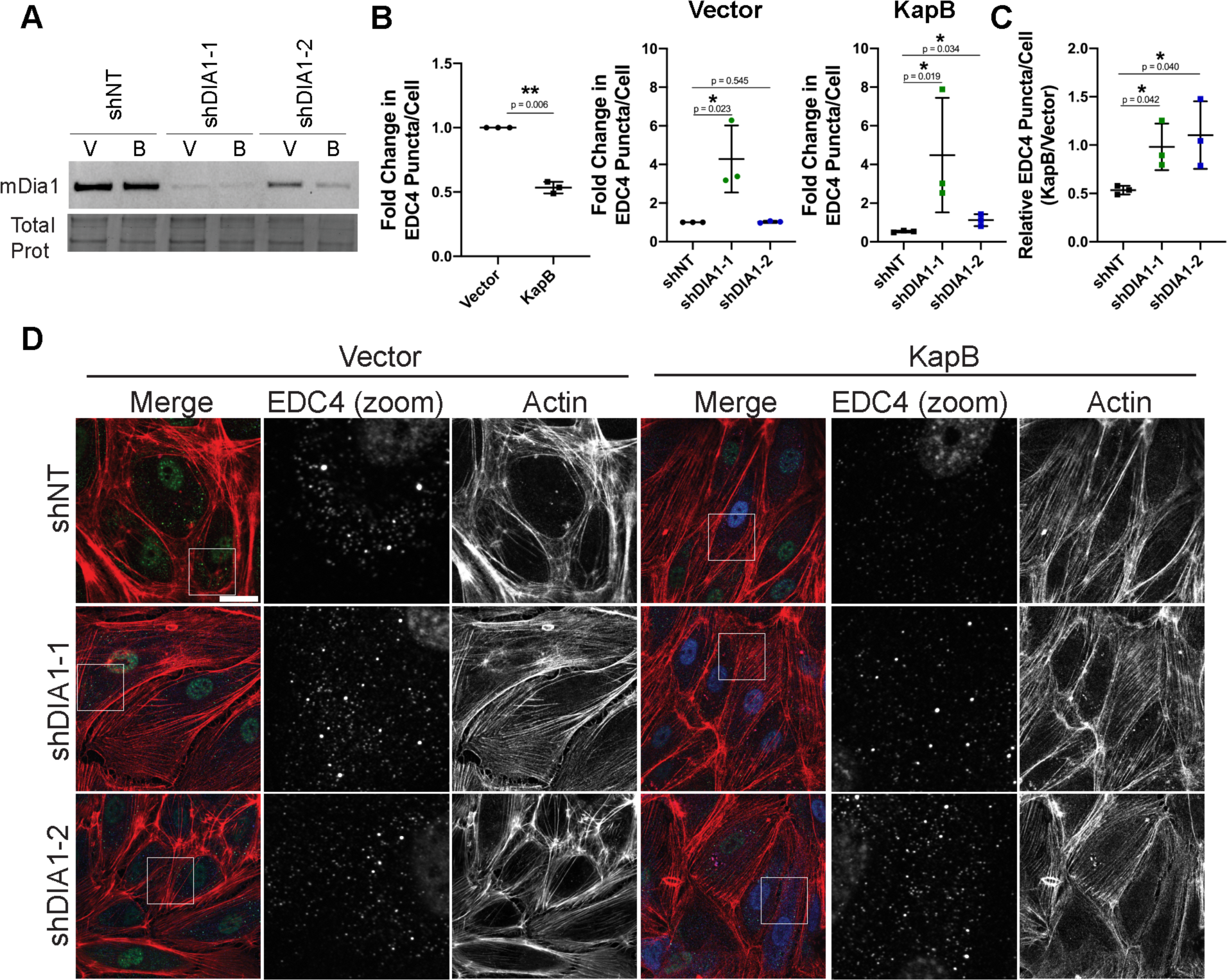
The RhoA-effector mDia1 is required for KapB-mediated PB disassembly. KapB- and vector- expressing HUVECs were transduced with shRNAs targeting mDia1 (shDIA1-1, shDIA1-2) or with a non-targeting (shNT) control and selected. In parallel, cells were fixed for immunofluorescence or lysed for immunoblotting. (A) One representative immunoblot of three independent experiments stained with mDia1-specific antibody is shown. (B, C) Fixed cells were stained for CellProfiler analysis as detailed in the methods. (B) The number of EDC4 puncta per cell was quantified and normalized to the vector NT control within each replicate. (C) CellProfiler data was used to calculate the ratio of EDC4 puncta counts in KapB-expressing cells versus the vector control for each treatment condition. (D) Representative images of cells stained for PB-resident protein EDC4 (green), KapB (blue), and F-actin (red, phalloidin). Boxes indicate area shown in the EDC4 (zoom) panel. Scale bar represents 20 µm. Statistics were determined using ratio paired t-tests between control and experimental groups; error bars represent standard deviation; n=3 independent biological replicates; * = p < 0.05, ** = p < 0.01.

ROCK promotes SF formation by increasing actin contractility and inhibiting actin severing activity (Julian and Olson 2014). Chemical inhibition of both isoforms of ROCK, ROCK1 and ROCK2, with Y-27632 (Ishizaki et al. 2000) restored PBs in KapB-expressing cells and increased the ratio of KapB:Vector PBs (Fig 2A-C). HUVECs treated with Y-27632 had scalloped cell edges with minimal actin structure (Fig 2A). Y-27632 did not visibly alter cell viability during the indicated treatment. To determine whether PB disassembly is dependent on a single ROCK isoform, both ROCK1 and ROCK2 were knocked down with isoform-specific shRNAs. Knockdown efficacy was confirmed with immunoblotting (Fig S1). Independent knockdown of ROCK1 and 2 increased PBs counts in KapB-expressing cells (Fig 2D, F) and restored the ratio of KapB:Vector PBs counts (Fig 2E). This indicated that both ROCK1 and ROCK2 can contribute to KapB-mediated PB disassembly. ROCK2 knockdown showed more robust PB restoration, both in terms of PB counts and PB size, than that seen with ROCK1 knockdown (Fig 2D, F). Quantification of PB counts in control cells for both pan-ROCK inhibition and ROCK knockdown experiments is reported in Figure S1. While pan-ROCK inhibition and ROCK1 knockdown treatments both eliminate SFs, ROCK2 knockdown retains pronounced actin fibres in the cells (Fig 2F). Similar to mDia1 knockdown, this may indicate a compensatory mechanism to retain cell shape and suggests that only a subset of SFs may be required for PB disassembly. Taken together, these data show that inhibition of RhoA effectors that mediate SF formation can reverse KapB-mediated PB disassembly. Put another way, we have been unable to uncouple KapB-mediated inducement of SFs from KapB-mediated PB disassembly.

**Figure 2:**
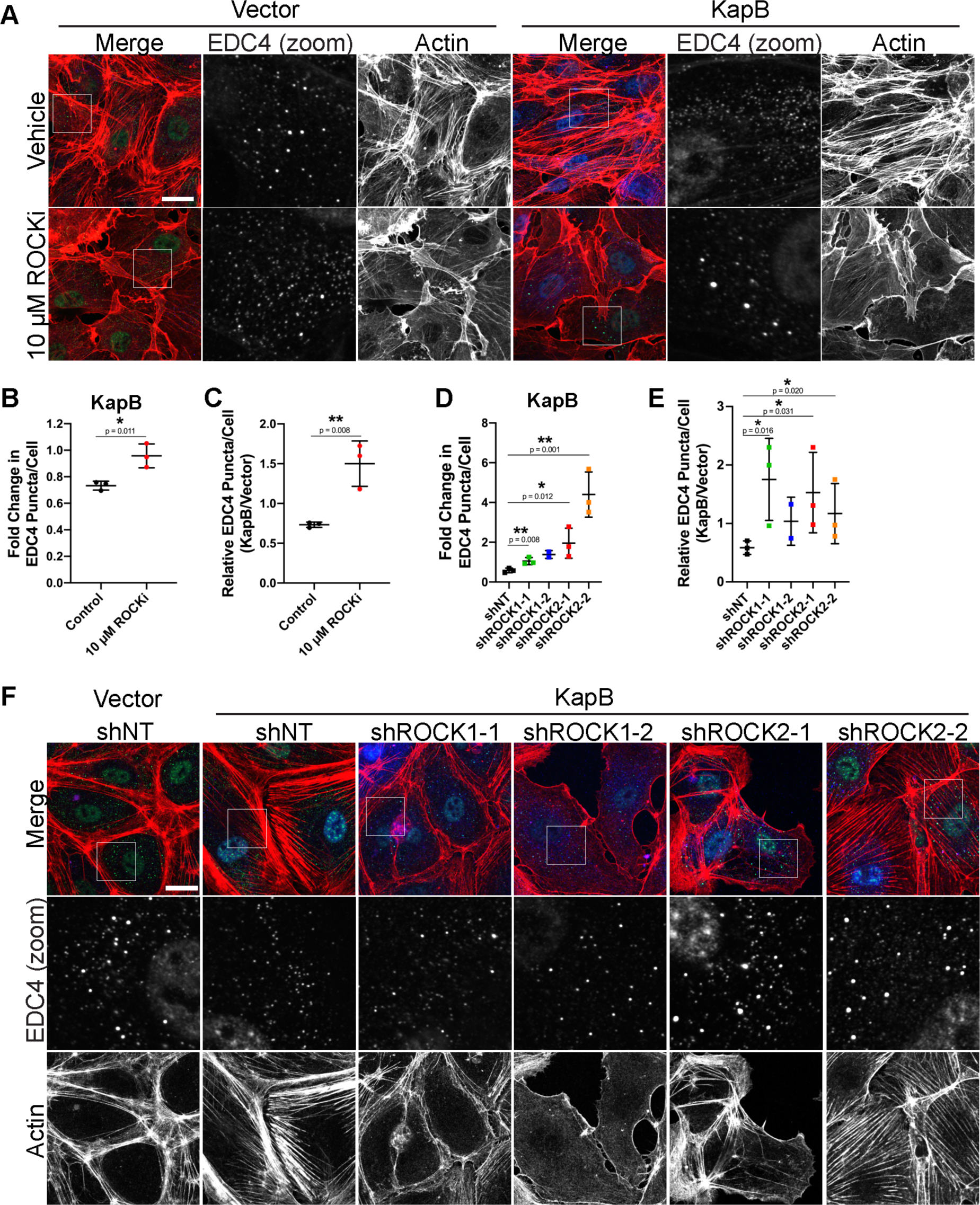
The RhoA-effector ROCK is required for KapB-mediated PB disassembly. (A-C) KapB- and vector- expressing HUVECs were treated with 10 μM Y-27632 or water control for 4 h and fixed for immunofluorescence. (A) Representative images of cells stained for PB-resident protein EDC4 (green), KapB (blue), and F-actin (red, phalloidin). Boxes indicate area shown in the EDC4 (zoom) panel. Scale bar represents 20 µm. (B, C) Fixed cells were stained for CellProfiler analysis as detailed in the methods. (B) The number of EDC4 puncta per cell was quantified and normalized to the vector NT control within each replicate. (C) CellProfiler data was used to calculate the ratio of EDC4 puncta counts in KapB-expressing cells versus the vector control for each treatment condition. (D-F) KapB- and vector- expressing HUVECs were transduced with shRNAs targeting ROCK1 and ROCK2 (shROCK1-1, shROCK1-2, shROCK2- 1, shROCK2-2) or with a non-targeting (shNT) control and selected. Cells were fixed for immunofluorescence. (D, E) Fixed cells were stained for CellProfiler analysis as detailed in the methods. (D) The number of EDC4 puncta per cell was quantified and normalized to the vector NT control within each replicate. (E) CellProfiler data was used to calculate the ratio of EDC4 puncta counts in KapB-expressing cells versus the vector control for each treatment condition. (F) Representative images of cells stained for PB-resident protein EDC4 (green), KapB (blue), and F-actin (red, phalloidin). Boxes indicate images shown in EDC4 (zoom) panel. Scale bar represents 20 µm. Statistics were determined using ratio paired t-tests between control and experimental groups; error bars represent standard deviation from n=3 independent biological replicates except shROCK1-2, (n=2); * = p < 0.05, ** = p < 0.01.

ROCK phosphorylates and activates LimK, which then phosphorylates and inactivates cofilin, an actin-severing protein (Ohashi et al. 2000). In this way, ROCK promotes SF formation by inactivating cofilin. To investigate the role of cofilin in KapB-mediated PB disassembly, shRNAs to knockdown cofilin expression were used (Fig S2). Since ROCK activation results in less cofilin activity and reduced actin severing, we hypothesized that knockdown of cofilin in KapB-expressing cells would augment KapB-mediated PB disassembly. Knockdown of cofilin resulted in elongated cells with more SFs in both control and KapB-expressing cells (Fig S2). Cofilin knockdown also induced PB disassembly in control cells and aided PBs disassembly in KapB-expressing cells (Fig S2 B, C). This indicates that inhibition of cofilin elicits PB disassembly and supports the hypothesis that reducing cofilin activity is one of the reasons that KapB promotes SF formation and PB disassembly.

### Changes in cytoskeletal contractility control PB disassembly

One of the downstream activities of the kinase, ROCK, is to phosphorylate myosin light chain to induce non-muscle myosin II (NMII)-mediated actomyosin contraction (Mutsuki Amano et al. 1996). Since ROCK is required for KapB-mediated PB disassembly, we determined whether functional actomyosin contractility is also required. KapB-expressing cells were treated with blebbistatin, which inhibits NMII-mediated actomyosin contractility by maintaining NMII in a conformation that is unable to bind actin filaments (Kovacs et al. 2004). Treatment of KapB-expressing cells with blebbistatin restored both PBs levels in KapB-expressing cells, as well as the KapB:Vector ratio of PBs (Fig 3A-C), indicating that contractility is required for KapB-induced PB disassembly. ECs treated with blebbistatin had normal actin structure (Fig 3C). Blebbistatin treatment for the indicated time did not visibly alter cell viability. To determine if contraction would elicit the same phenotype in the absence of KapB, cells were treated with Calyculin A (CalA), an inhibitor of myosin light chain phosphatase that promotes NMII phosphorylation and actomyosin contraction (Asano and Mabuchi 2001). CalA treatment did not visibly alter actin cytoskeletal structure or cell viability. Inducing contraction with CalA decreased counts of PBs (Fig 3D, E), again consistent with the hypothesis that actomyosin contractility controls PB disassembly. Actomyosin contractility impacts cytoskeletal tension in adherent cells with SFs (Katoh et al. 1998; Tan et al. 2003). Since the mechanoresponsive transcription activator, YAP, is activated by increases to cytoskeletal tension via actomyosin contractility (Dupont et al. 2011), we explored the role of YAP in KapB-mediated PB disassembly.

**Figure 3:**
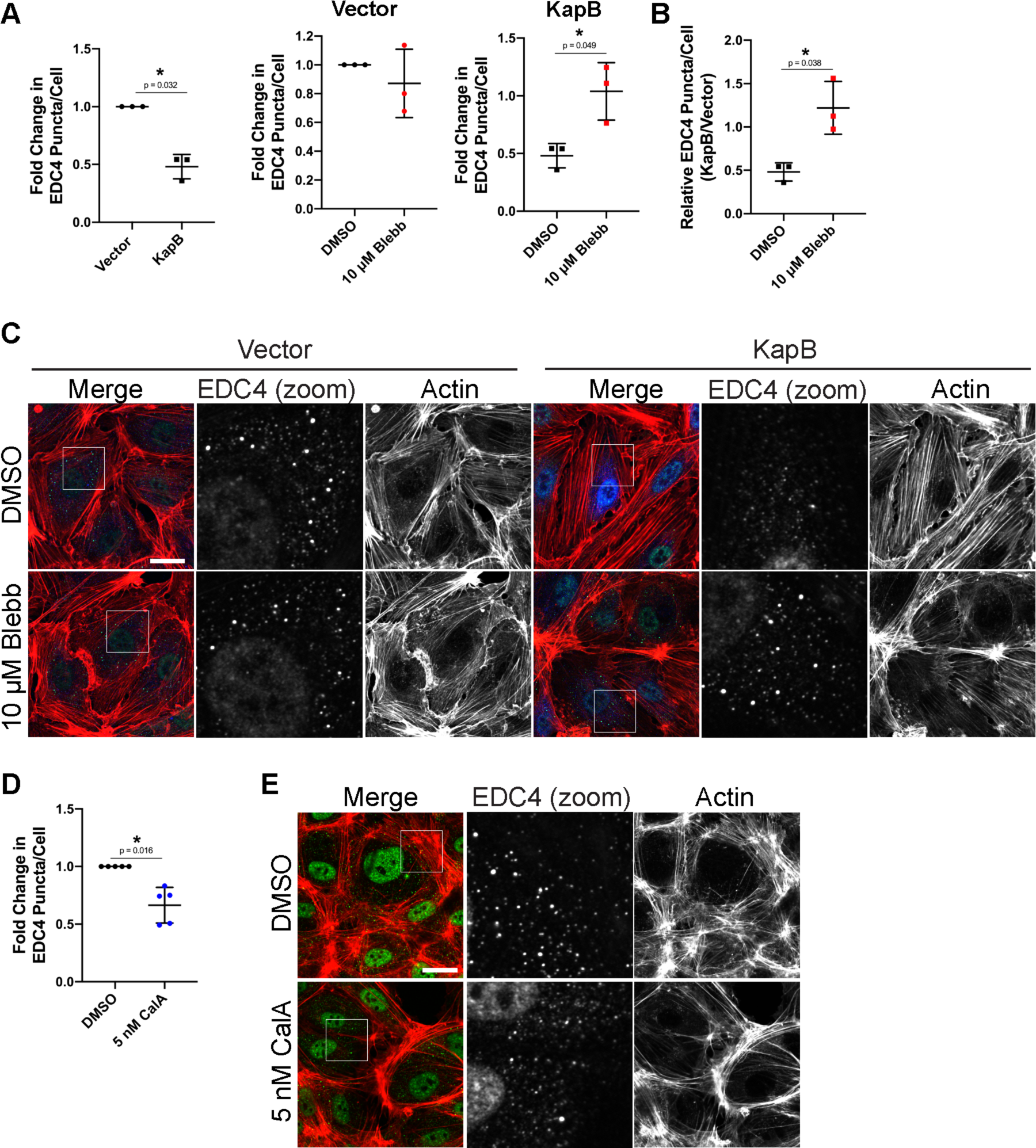
Actomyosin contractility controls PB disassembly. (A-C) KapB- and vector- expressing HUVECs were treated with 10μM blebbistatin to inhibit actomyosin contractility or DMSO for 30 min and fixed for immunofluorescence. (A, B) Fixed cells were stained for CellProfiler analysis as detailed in the methods. (A) The number of EDC4 puncta per cell was quantified and normalized to the vector NT control within each replicate. (B) CellProfiler data was used to calculate the ratio of EDC4 puncta counts in KapB-expressing cells versus the vector control for each treatment condition. (C) Representative images of cells stained for PB-resident protein EDC4 (green), KapB (blue), and F-actin (red, phalloidin). Boxes indicate area shown in the EDC4 (zoom) panel. Scale bar represents 20 µm. (D, E) Untransduced HUVECs were treated with 5 nM Calyculin A (CalA) to stimulate actomyosin contraction or DMSO for 20 min and fixed for immunofluorescence. (D) Fixed cells were stained for CellProfiler analysis as detailed in the methods. EDC4 puncta per cell were quantified and normalized to the DMSO control within each replicate. (E) Representative images of cells treated with 5 nM CalA and stained for PB-resident protein EDC4 (green) and F-actin (red, phalloidin). Boxes indicate area shown in the EDC4 (zoom) panel. Scale bar represents 20 µm. Statistics were determined using ratio paired t- tests between control and experimental groups; error bars represent standard deviation; n=3 (A, B) and n=5 (D) independent biological replicates; * = p < 0.05.

### YAP is required for KapB-mediated PB disassembly

To determine if YAP was involved in KapB-mediated PB disassembly, we expressed shRNAs targeting YAP in KapB-expressing HUVECs to assess whether the altered levels of YAP impacted PB disassembly. Immunoblotting confirmed knockdown efficiency of YAP (Fig 4A). Knockdown of YAP resulted in elongated HUVECs with mostly cortical actin fibres (Fig 4D). These cells displayed increased PBs in KapB-expressing cells (Fig 4B-D). In the context of YAP knockdown, the KapB:Vector ratio of PBs counts was restored, indicating that YAP is required for KapB-mediated PB disassembly (Fig 4C) and suggesting that KapB is activating a mechanoresponsive signalling axis to elicit PB disassembly via YAP.

**Figure 4:**
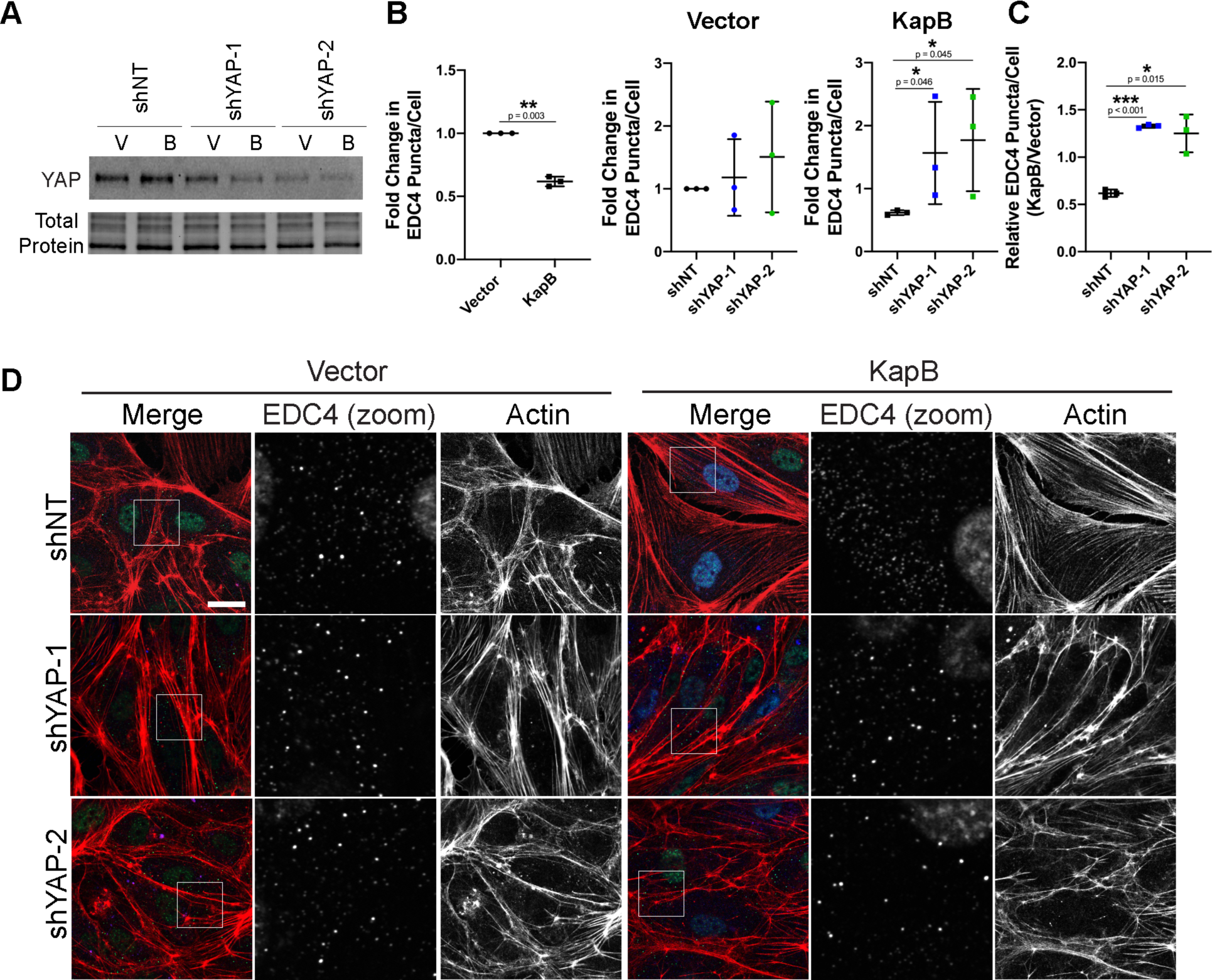
YAP is required for KapB-mediated PB disassembly. KapB- and vector- expressing HUVECs were transduced with shRNAs targeting YAP (shYAP-1, shYAP-2) or with a non-targeting (shNT) control and selected. In parallel, cells were fixed for immunofluorescence or lysed for immunoblotting. (A) One representative immunoblot of three independent experiments stained with YAP-specific antibody is shown. (B-D) Fixed cells were stained for CellProfiler analysis as detailed in the methods. (B) The number of EDC4 puncta per cell was quantified and normalized to the vector NT control within each replicate. (C) CellProfiler data was used to calculate the ratio of EDC4 puncta count in KapB-expressing cells to the vector control for each treatment condition. (D) Representative images of cells stained for PB-resident protein EDC4 (green), KapB (blue), and F-actin (red, phalloidin). Boxes indicate area shown in the EDC4 (zoom) panel. Scale bar represents 20 µm. Statistics were determined using ratio paired t-tests between control and experimental groups; error bars represent standard deviation; n=3 independent biological replicates; * = p < 0.05, ** = p < 0.01, *** = p < 0.001.

We next investigated the steady-state protein level and localization of YAP in KapB-expressing cells. KapB-transduced HUVECs showed increased levels of nuclear YAP and total YAP in some experiments, though the ratio of nuclear:cytoplasmic YAP was not markedly increased (Fig S3A-B). When YAP is phosphorylated by LATS, most notably at serine 127, it is sequestered in the cytoplasm and transcriptionally inactive (Zhao et al. 2007). To investigate the phosphorylation status of YAP in KapB-expressing cells, levels of P(S127)-YAP and total YAP were measured by immunoblot. In KapB-expressing cells, there was a decrease in the ratio of P(S127)-YAP to total YAP that may suggest YAP is more active when KapB is expressed (Fig S3B). In some experiments, we observed an increase in total steady-state levels of YAP by immunoblotting, corroborating the small increase in total YAP intensity seen by microscopy (Fig S3A-B). Although increased total YAP may suggest that YAP turnover is blocked by KapB, these increases were not accompanied by a strong decrease in phosphorylated YAP or an increase in nuclear YAP, as in (Pavel et al. 2018).

We next asked if active YAP can interact with TEAD and other transcription factors to elicit changes in gene expression in KapB-expressing cells (Vassilev et al. 2001). We used YAP 5SA, a constitutively active version of YAP that is unable to be phosphorylated and inactivated by the inhibitory kinase LATS, as a positive control (Zhao et al. 2007). KapB did not increase steady-state mRNA levels of common YAP target genes CTGF, CYR61 and ANKRD1 by RT-qPCR, although these genes were elevated by YAP 5SA (Fig S3C). These data indicate despite the observation that YAP may be more abundant in KapB-expressing cells, canonical YAP targets are not upregulated. We wondered if YAP activation could elicit PB disassembly in the absence of KapB expression. To this end, we transduced HUVECs with constitutively active YAP 5SA. YAP 5SA-expressing cells contained few, thick actin fibres (Fig 5A) and displayed decreased number of PBs per cell, indicating that YAP 5SA elicited disassembly of PBs (Fig 5A, B). We also confirmed that we could stain PBs in control cells with antibodies for more than one PB resident protein (EDC4, Dcp1a and DDX6) and that when YAP 5SA was expressed, loss of PB puncta was observed for each of these staining conditions (Fig 5 A-E). Although PB disassembly is induced by the expression of both YAP 5SA and KapB, active YAP increases the transcriptional activation of genes CTGF, CYR61 and ANKRD1 while KapB does not.

**Figure 5:**
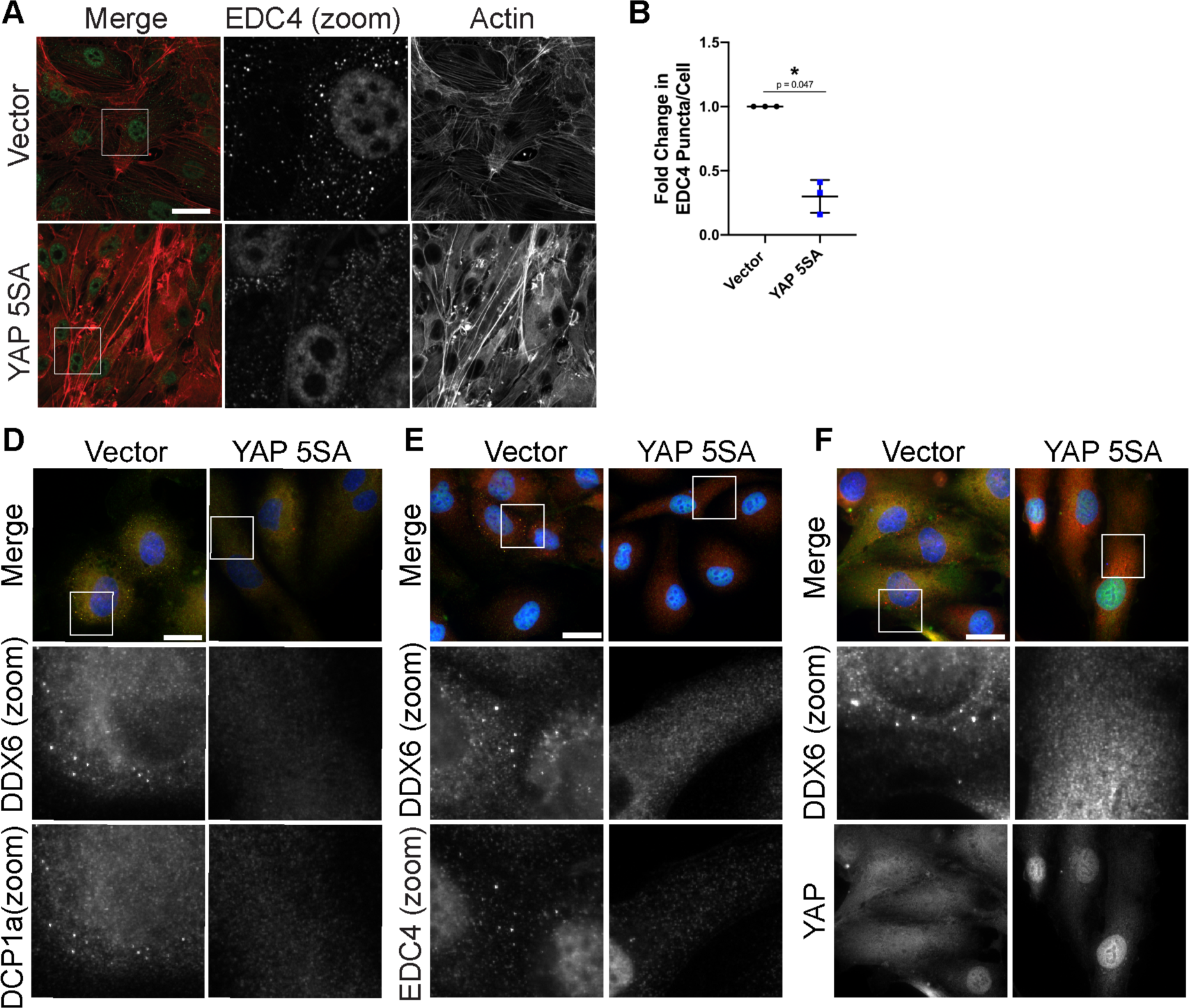
Active YAP elicits PB disassembly. HUVECs were transduced with YAP 5SA- expressing and empty vector lentivirus and selected. Cells were fixed for immunofluorescence. (A) Representative images of cells stained for PB-resident protein EDC4 (green) and F-actin (red, phalloidin). Boxes indicate area shown in the EDC4 (zoom) panel. Scale bar represents 20 µm. (B) Fixed cells were stained for CellProfiler analysis as detailed in the methods. The number of EDC4 puncta per cell was quantified and normalized to the vector control. (C-E) Representative images of cells stained for (C) DDX6 (red), DCP1a (green) and DAPI (blue), (D) DDX6 (red), EDC4 (green) and DAPI (blue), and (E) DDX6 (red), YAP (green) and DAPI (blue). Boxes indicate area shown in the zoom panels. Scale bar represents 20 µm. Statistics were determined using ratio paired t-tests between control and experimental groups (B); error bars represent standard deviation; n=3 independent biological replicates; * = p < 0.05, ** = p < 0.01.

We wondered if PB disassembly did depend on the ability of YAP to transactivate transcription of its canonical genes. To better understand this, we transduced HUVECs with another YAP construct called YAP 6SA (Fig 6). YAP 6SA contains the same mutations as 5SA; in addition, it cannot be phosphorylated by AMPK at serine 94 (Mo et al. 2015). This phosphorylation event is essential for its interaction with TEAD; therefore, the S94A mutation renders YAP 6SA transcriptionally inactive (Mo et al. 2015, Zhang et al. 2017). We observed that unlike YAP 5SA or KapB, YAP 6SA did not appear to elongate cells, suggesting that actin dynamics were unaffected, as in Pavel et al. (2018). Compared to YAP 5SA which caused pronounced PB disassembly, YAP 6SA failed to alter PB levels compared to vector controls (Fig 6B). This was confirmed by staining PBs with antibodies for two different resident proteins, DDX6 and EDC4 (Fig 6A).

**Figure 6:**
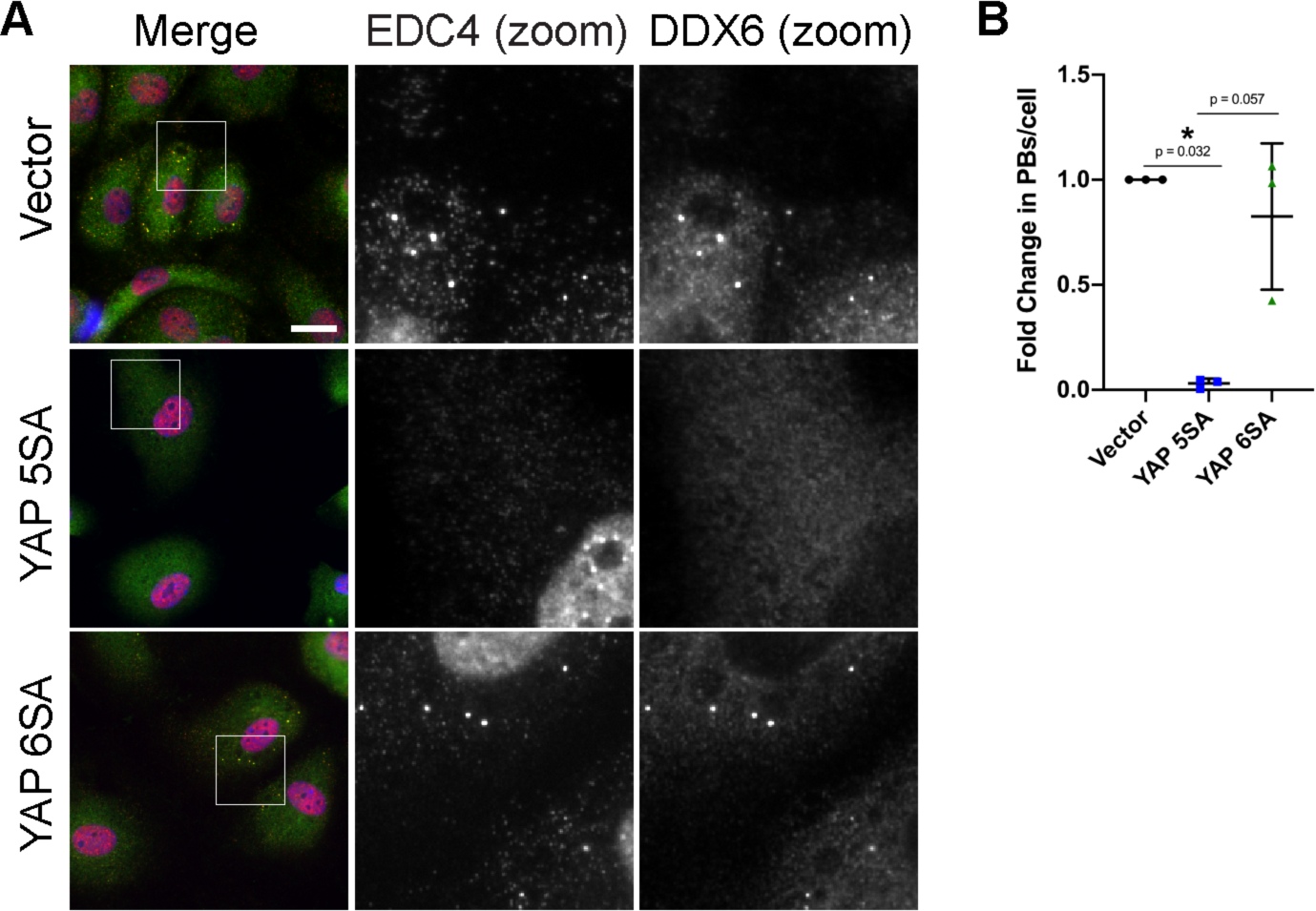
YAP transcriptional activity is required for PB disassembly. HUVECs were transduced with YAP 5SA-, YAP 6SA- or vector-expressing lentivirus and selected. Cells were fixed for immunofluorescence. (A) Representative images of cells stained for EDC4 (red), DDX6 (green) and DAPI (blue). Boxes indicate area shown in the zoom panels. Scale bar represents 20 µm. (B) Fixed cells were stained for CellProfiler analysis as detailed in the methods. CellProfiler was used to count nuclei, EDC4 puncta and DDX6 puncta. In RStudio analysis, puncta with ≥ 60% correlation (mean correlation in vector control) between EDC4 and DDX6 (PBs) were counted and normalized to number of nuclei per condition. PB counts were normalized to vector control within each replicate.

### YAP activators disassemble PBs

Since overexpression of constitutively active YAP 5SA leads to disassembly of PBs, we wanted to determine whether other stimuli that activated endogenous YAP could do the same. We tested various upstream mechanical signals described to activate YAP for their ability to elicit PB disassembly: shear stress, low cell confluence and high ECM stiffness (Nakajima et al. 2017; Lee et al. 2017; Noria et al. 2004; Zhao et al. 2007; Dupont et al. 2011). For the first, we subjected HUVECs to shear stress by fluid flow (shear forces of 2 and 10 dyn/cm^2^) and PBs were examined via immunofluorescence using antibodies to both EDC4 and DDX6. Both treatments showed prominent cell elongation, stress fibre formation, and resulted in robust PB disassembly of both EDC4-positive and DDX6-positive puncta (Fig 7A, B). To test if cell confluence regulates PB levels, HUVECs were seeded at low, medium and high densities. Cells at low confluence are reported to have active YAP and we predicted PBs would disassemble; however, the low-density monolayer displayed more PBs per cell then those at medium and high densities (Fig 7C, D). To test the impact of collagen stiffness on PB disassembly, HUVECs were plated on coverslips coated with increasing densities of collagen (0 to 64 µg/ cm^2^). While collagen density does not perfectly reproduce matrix stiffness as it does not differentiate the effect of matrix stiffness from increasing collagen-binding sites, increasing collagen densities do correlate with increased matrix stiffness (Yang, Leone, and Kaufman 2009; Lee et al. 2014; Joshi, Mahajan, and Kothapalli 2018). As collagen density increased, PBs decreased (Fig 7E, F). Notably, as collagen density increased, only slight increases in visible actin fibres were noted (Fig 7E), suggesting the tested range of stiffness was small and should be expanded in future studies. Taken together, these data indicate that PB disassembly occurred in response to mechanical stimuli known to require RhoA and altered cytoskeletal structures to activate YAP (shear stress and increased ECM concentration) (Zhao et al. 2012; Huang et al. 2016; Lee and Kumar 2016; Moreno-Vicente et al. 2018) but not in response to YAP stimuli that do not also change actin cytoskeletal structures (low cell confluence). These data support the importance of actin SF formation and YAP activation as requisite precursors for PB disassembly not only YAP activation status.

**Figure 7:**
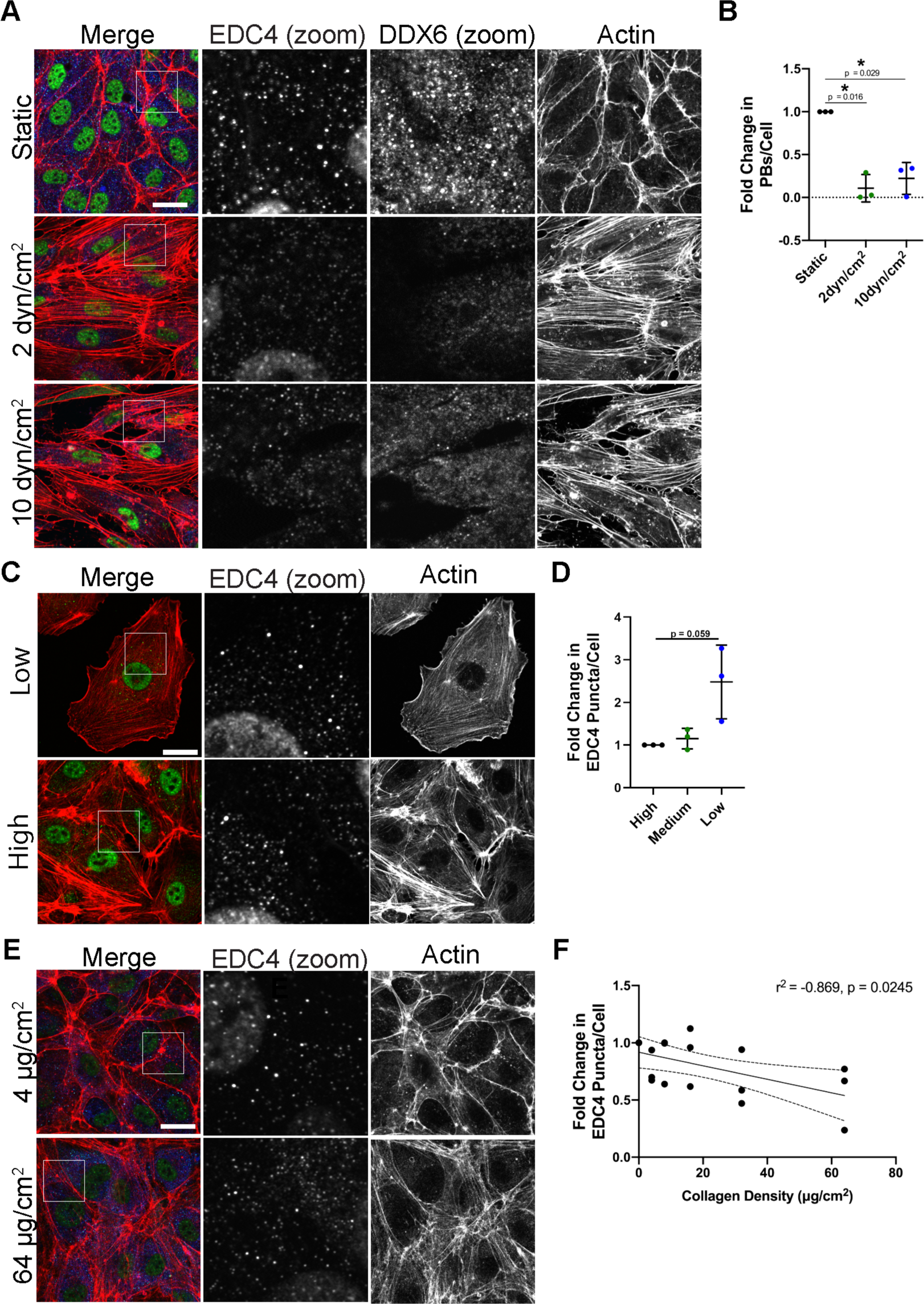
YAP Inputs Mediate PB Disassembly. (A, B) HUVECs were seeded onto collagen- coated microscope slides and exposed to shear stress of 2 dyn/cm^2^, 10 dyn/cm^2^ or no shear (static control) for 21 h. Cells were fixed and stained for immunofluorescence. (A) Representative images of cells stained for PB-resident proteins EDC4 (green) and DDX6 (blue), as well as F- actin (red, phalloidin). Boxes indicate area shown in EDC4 (zoom) and DDX6 (zoom) panels. Scale bar represents 20 µm. (B) CellProfiler was used to count nuclei, EDC4 puncta and DDX6 puncta. In RStudio analysis, puncta with ≥ 70% correlation (mean correlation in vector control) between EDC4 and DDX6 (PBs) were counted and normalized to number of nuclei per condition. PB counts were normalized to static control within each replicate. (C, D) HUVECs were split and seeded at a high-, medium- and low-density, cultured for 48 h and fixed for immunofluorescence. (C) Representative images of cells stained for the PB-resident protein EDC4 (green) and F-actin (red, phalloidin). Boxes indicate images shown in EDC4 (zoom) panel. Scale bar represents 20 µm. (D) Fixed cells were stained for CellProfiler analysis as detailed in the methods. The number of EDC4 puncta per cell was quantified and normalized to the high confluence condition. (E, F) Coverslips were coated for 4 h with 0 to 64 µg/cm^2^ of collagen. HUVECs were grown for 72 h on coated coverslips and fixed for immunofluorescence. (E) Representative images of cells stained for PB-resident protein EDC4 (green), DDX6 (blue) and F-actin (red, phalloidin). Boxes indicate images shown in EDC4 (zoom) panel. Scale bar represents 20 µm. (F) Fixed cells were stained for CellProfiler analysis as detailed in the methods. The number of EDC4 puncta per cell was quantified and normalized to the 0 µg/mL collagen-coating condition. Statistics were determined using repeated measures ANOVA (A, B) or Pearson’s correlation co-efficient (C); error bars represent standard deviation (A, B) and 95% confidence interval of line of best fit (slope is significantly non-zero, p = 0.014) (C); n=3 independent biological replicates; * = p < 0.05, ** = p < 0.01.

### Shear stress mediated PB disassembly requires YAP

YAP responds to external forces that induce active RhoA, actin SFs, and pronounced cell elongation; in short, the typical behaviour of ECs in response to the mechanical shear stress that is produced by fluid flow. However, how YAP responds to shear stress is less clear (Wang et al. 2016; Huang et al. 2016; Lee et al. 2017; Nakajima and Mochizuki 2017). To verify YAP activation by continuous, unidirectional fluid flow in our system, HUVECs subjected to 2 and 10 dyn/cm^2^ of shear stress were lysed and used for immunoblotting for P(S127)-YAP and total YAP. Shear stress decreased the ratio of phospho-YAP/YAP in both conditions, suggesting a higher proportion of active YAP (Fig 8A). To assess if YAP was required for PB disassembly in response to shear stress, HUVECs transduced with YAP-targeting shRNA were subjected to 10 dyn/cm^2^ shear stress. PBs disassembled in cells treated with a non-targeting shRNA when subjected to shear stress (Fig 8B, C), consistent with earlier experiments (Fig 7A, B). When YAP was reduced by shRNA expression, ECs exposed to shear stress had more PBs than static growth control cells (Fig 8B, C). YAP knockdown also reduced the cells ability to form SFs, with many cells displaying highly cortical phalloidin staining and fewer actin fibres across the cell (Fig 8C). Therefore, YAP is required to disassemble PBs after KapB is expressed and also in response to shear stress.

**Figure 8:**
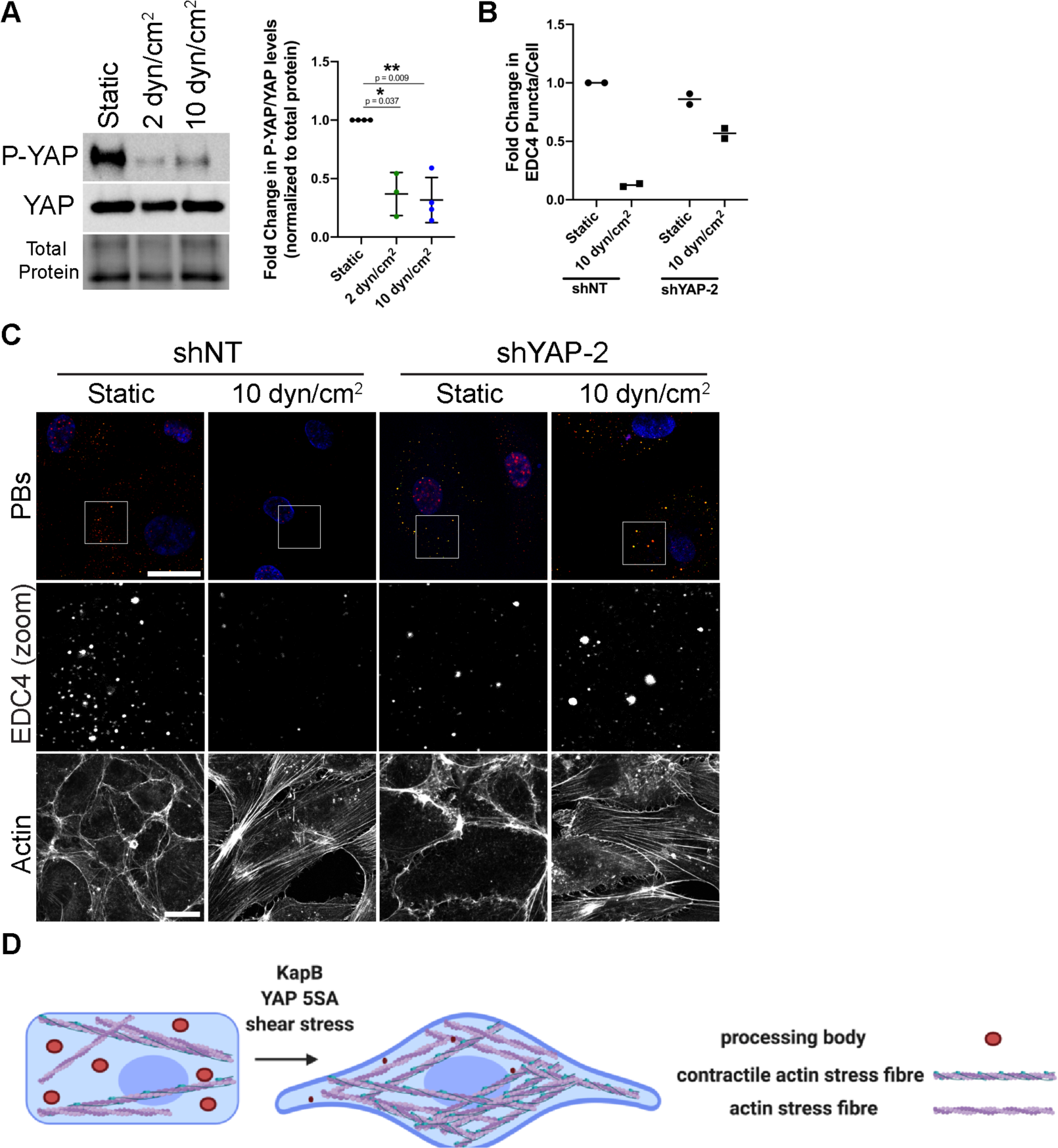
YAP is required for EDC4 puncta disassembly in HUVECs subjected to shear stress. (A) HUVECs were seeded onto collagen-coated microscope slides and exposed to shear stress of 2 dyn/cm^2^, 10 dyn/cm^2^ or a static control for 21 h. Cells were lysed for immunoblotting. One representative immunoblot and quantification of three independent experiments stained with P(S127)-YAP- and YAP-specific antibody is shown. P(S127)-YAP and YAP protein levels in each condition were normalized to total protein. All treatments were normalized to static control within each replicate. (B, C) HUVECs were transduced with shRNAs targeting YAP (shYAP-2) or with a non-targeting (shNT) control and selected. Cells were seeded onto collagen-coated microscope slides and exposed to shear stress of 10 dyn/cm^2^ or no shear (static control) for 21 h. Cells were fixed and stained for immunofluorescence. (B) CellProfiler was used to count nuclei and EDC4 puncta. In RStudio analysis, EDC4 puncta were normalized to number of nuclei per condition. EDC4 puncta counts were normalized to static control. (C) Representative images of cells stained for PB-resident protein EDC4 (red), DDX6 (green) and DAPI (blue). In parallel, separate coverslips were stained for F-actin (phalloidin). Boxes indicate area shown in the EDC4 (zoom) panel. Scale bar represents 20 µm. Statistics were determined using repeated measures ANOVA (A); error bars represent standard deviation (A) ; n=4, except 2 dyn/cm^2^ (n=3) (A) and n = 2 (B, C) independent biological replicates; * = p < 0.05, ** = p < 0.01. (D) KapB activates a mechanoresponsive pathway from within the cell rather than without to mediate PB disassembly. Cells respond to external mechanical force by activating their structural support network, the actin cytoskeleton. The GTPase RhoA and its downstream effectors coordinate this response, bundling actin filaments into stress fibers (SFs), enhancing actomyosin contractility and increasing adhesion to the underlying matrix to help withstand force-induced membrane deformation. Together, these actin-based responses increase cytoskeletal tension and elicit the dephosphorylation and nuclear translocation of the mechanoresponsive transcription activator YAP where it collaborates with other transcription factors to induce TEAD-responsive genes. We present data to support the existence of a mechanoresponsive pathway that links actin SFs, actomyosin contractility, and the transcription transactivator YAP to the disassembly of PBs. The viral protein KapB taps into this mechanoresponsive pathway, triggering mechanical changes and forming contractile cytoskeletal structures that would normally respond to force, thereby inducing PB disassembly in a YAP-dependent manner, from within the cell, rather than from without. Both KapB and stimuli that activate YAP cause PB disassembly.

Taken together with our analysis of KapB-mediated PB disassembly, these data suggest that when KapB is expressed, it turns on mechanoresponsive signals that cells use to withstand mechanical forces like shear, in the absence of an external stimulus. The outcome of both KapB expression and exposure to mechanical forces is YAP-dependent disassembly of cytoplasmic PBs (Fig 8D).

## Discussion

In this manuscript, we have used a viral protein from an oncogenic virus to uncover a new relationship between cytoplasmic PBs and the mechanical regulation of actin SF formation and YAP. We present data to support the existence of a mechanoresponsive pathway that links actin SFs, actomyosin contractility, and the transcription transactivator YAP to the disassembly of PBs and show that this pathway is hijacked by KapB during KSHV latency. Our major findings are as follows. i) KapB-mediated PB disassembly requires actin SF effectors ROCK1/2 /mDia1 and is enhanced by loss of the actin-severing protein, cofilin. ii) KapB-mediated PB disassembly is reversed when blebbistatin is used to inhibit actomyosin contractility or after knockdown of the mechanoresponsive transcription transactivator, YAP. iii) In the absence of KapB, we can induce PB disassembly when we promote actomyosin contractility and YAP activity using Calyculin A (inhibits myosin light chain phosphatase to promote actomyosin contraction (Asano and Mabuchi 2001)), or overexpression of active YAP (YAP 5SA). Exposure of endothelial cells to mechanical forces created by shear stress or a stiff extracellular matrix also induces PB disassembly in the absence of KapB. iv) Overexpression of transcriptionally inactive YAP (YAP 6SA) fails to disassemble PBs, indicating that YAP’s role as a transcription transactivator is important for its ability to promote PB disassembly. Together, these data show for the first time, that PBs disassemble in response to mechanical signals that transduce external forces from outside the cell to the actin cytoskeleton and that this pathway requires YAP. Moreover, this work also highlights the remarkable pizzazz used by viruses to hijack cellular pathways. In this case, we reveal that the viral protein KapB taps into this mechanoresponsive pathway to trigger mechanical changes to cytoskeletal structures and downstream effectors like YAP that would normally respond to force, thereby inducing PB disassembly from within the cell, rather than from without.

Although SFs with periodic distribution of actin-crosslinking proteins and non-muscle myosin II (NMII) have the potential to be contractile structures, in order to be able to generate tension, SFs must be tethered at the ends (Katoh et al. 1998; Tan et al. 2003). Ventral SFs are attached at both termini to the extracellular matrix (ECM) through focal adhesions and contain NMII, which imparts a contractile phenotype, whereas dorsal SFs are attached through focal adhesions but do not contain NMII, and thus are not contractile (Hotulainen and Lappalainen 2006; Small et al. 1998; Vallenius 2013). Several features of our data suggest that the actin structures that are important for KapB-mediated PB disassembly must be contractile and cause cytoskeletal tension. When both mDia1 and ROCK2 were silenced in KapB-expressing cells (Fig 1, 2), visible actin bundles are still apparent despite PB restoration in both contexts. This suggests that not all SF subtypes are required for our phenotype. In addition, blebbistatin treatment of KapB-expressing cells dramatically restored PBs; these data suggest that PB disassembly requires actin-mediated contractility rather than structural support (Fig 3). Furthermore, shear stress and increased ECM stiffness (Lee et al. 2006; Jackson et al. 2008) both induced PB disassembly (Fig 7), reinforcing the correlation between increasing cell tension and PB disassembly. Finally, our data show that YAP is required for PB disassembly (Figs 4, 5, 8). YAP is mechanoresponsive; it becomes active when tension-forming actin structures are induced by external forces e.g. focal adhesion engagement by stiff ECM (Dupont et al. 2011; Sugimoto et al. 2018). As YAP activation and PB disassembly both rely on RhoA-induced cytoskeletal contractility, any activator of YAP that induces cytoskeletal tension through RhoA should mediate PB disassembly. Our data support this notion, as shear stress forces and increasing collagen density both cause PB disassembly in the absence of KapB, while low confluence does not (Fig 7). We also know that GPCRs (G_11/12_ and G_q/_11) activate YAP in a RhoA-dependent manner (Yu et al. 2012) and LPA treatment or overexpression of KSHV-derived constitutively active vGPCR (both activate G_11/12_) both induce PB disassembly (Corcoran et al. 2012; Corcoran and McCormick 2015). These findings support the conclusion that PB disassembly requires the formation of contractile actin structures like those associated with YAP transactivation responses.

Data presented herein clearly implicate a requirement for YAP in the PB disassembly phenotype that is induced by KapB and by the external force imparted by shear stress (Fig 4, 8). However, the precise connection between YAP and PB disassembly is unclear. Despite the clear reliance on YAP for PB disassembly, KapB does not increase expression of canonical YAP-regulated transcripts (Fig S3). Our data also show small increases in total YAP and small decreases in the ratio of phosphorylated/total YAP; however, the ratio of nuclear:cytoplasmic YAP is not markedly altered (Fig S3). Taken together, these data first suggested a model whereby PB disassembly was independent of the role of YAP as a gene transactivator and YAP nuclear translocation. However, we found that overexpression of YAP 6SA, which is unable to act as a transcription transactivator, failed to cause PB disassembly (Fig 6), indicating that the transcription induction function of YAP is required for its role in regulating PB dynamics. Future experimentation is needed to further define the mechanistic link between YAP activation and PB disassembly.

KSHV is an oncogenic virus associated with the endothelial neoplasm, Kaposi’s sarcoma (KS). Cells within the KS lesion display latent KSHV infection, proliferate abnormally, spindle, and release many pro-inflammatory and pro-tumourigenic mediators into the microenvironment. KapB expression is sufficient to recapitulate two of these key features, cell spindling and pro-inflammatory mediator production that results from enhanced stability of ARE-containing cytokine mRNAs that would normally shuttle to PBs for constitutive turnover (Corcoran, Johnston, and McCormick 2015). Our previous work showed that both phenotypes utilize KapB activation of the stress-responsive kinase, MK2, and the downstream activation of the GTPase RhoA (Corcoran, Johnston, and McCormick 2015; Corcoran and McCormick 2015). We have now extended these studies to show that PB disassembly depends on mechanical signals that cause cell elongation, actin contractility and activate YAP. However, other KSHV gene products have been observed to induce cell elongation and/or cytoskeletal changes. These include the latent protein vFLIP, and two lytic proteins, vGPCR and the viral thymidine kinase (TK) (Corcoran et al. 2012, Grossman et al. 2006; Gill et al. 2005; Gill et al. 2015; Lagunoff 2015; Shepard et al. 2001). Both vFLIP and vGPCR cause cell elongation, while KSHV-TK causes RhoA-dependent cell contraction that leads to cell detachment (Corcoran et al. 2012, Grossman et al. 2006; Gill et al. 2015, Shepard et al. 2001). We previously showed that the lytic vGPCR protein mediates PB disassembly and the concomitant stabilization of ARE-mRNAs, (Corcoran et al. 2012); however, how vFLIP or KSHV-TK expression influences PB dynamics is unknown. The observation that KSHV encodes multiple gene products that alter cytoskeletal dynamics and promote inflammatory gene expression via PB disassembly further supports the linkage of these two phenotypes during KSHV infection.

PBs are sites where innate immune factors congregate that are disrupted by most viruses during infection (Burdick et al. 2010; Li et al. 2012; Ostareck, Naarmann-de Vries, and Ostareck-Lederer 2014; Burgess and Mohr 2015; Cuevas et al. 2016; H. Wang et al. 2016; Lumb et al. 2017; Balinsky et al. 2017; Núñez et al. 2018; Ng et al. 2020). KSHV encodes three separate proteins that all induce PB disassembly, KapB, vGPCR and ORF57, suggesting this event is central to viral persistence (Corcoran et al. 2012; Corcoran, Johnston, and McCormick 2015; Sharma et al. 2019). PBs are likely playing an as yet undefined and underappreciated role in regulating innate antiviral responses. KSHV encodes two proteins that influence YAP activity, suggesting YAP activation is also central to KSHV biology (Liu et al. 2015). YAP is an unappreciated negative regulator of innate immune signaling pathways. YAP blocks the ability of the innate immune kinase, TBK1, a downstream effector for several innate signaling pathways, to associate and activate its substrates (Zhang et al 2017). In so doing, YAP blocks downstream induction of interferons and increases viral replication (Zhang et al. 2017). However, this feature of YAP is independent of its ability to act as a transcriptional transactivator (Zhang et al. 2017). Based on our data that show YAP 5SA causes PB disassembly while YAP 6SA does not, KapB-induced PB disassembly requires the transcription transactivation function of YAP. We speculate that KapB-induced PB disassembly, like active YAP, favours viral replication and survival and is promoted by KSHV in order to reshape subsequent antiviral innate immune responses.

In this manuscript, we describe the surprising convergence of two previously unrelated yet essential regulators of cellular gene expression – the mechanoresponsive transactivator YAP and cytoplasmic PBs, known sites of AU-rich cytokine mRNA decay or translational suppression. We show that PB disassembly is mechanoresponsive; external forces that change cell shape and tension-forming cytoskeletal structures cause PB disassembly in a YAP-dependent manner. This discovery was made courtesy of the unique KSHV protein, KapB, and provides yet another example of how viruses have evolved surprising ways to manipulate their host and ensure their survival. In this case, KapB induces, from the inside out, a mechanoresponsive pathway to cause PB disassembly. Future study will untangle how these related mechanoresponsive events are induced by KSHV to better promote viral replication.

## Materials and Methods

### Antibodies, Plasmids and Reagents

The antibodies used in this study can be found in Table S1. The plasmids used in this study can be found in Table S2. Forward and reverse shRNA sequences were selected from the TRC Library Database in the Broad Institute RNAi consortium. YAP target shRNAs in pLKO.1 were obtained from Dr. C. McCormick (Dalhousie University, Halifax, Canada). Sequences for all shRNA oligonucleotides used for cloning are listed in Table S3. Cloning of shRNAs was conducted according to the pLKO.1 protocol (Addgene 2006). YAP 6SA was subcloned into pLJM1 by PCR amplification using primers (For: 5’-CGTAACCGGTATGGATCCCGGGCA- 3’, Rev: 5’-CTGATAAGTCGACAACCACTTTGTACAAGAAAGTTG-3’) designed to remove the V5 tag and insert a stop codon as well as 5’ AgeI and 3’ SalI restriction enzyme sites. The chemical inhibitors used in this study can be found in Table S4.

### Cell Culture

Human embryonic kidney 293T and 293A cells (HEK-293T/A, ATCC, Manassas, Virginia, US) and human cervical adenocarcinoma cells expressing a tetracycline-inducible repressor (HeLa Tet-Off, Clontech, Mountain View, California, US) were cultured in Dulbecco’s Modified Eagle Medium (DMEM, Gibco, Carlsbad, California, US) supplemented with 10% heat-inactivated fetal bovine serum (Gibco), 100 U/mL penicillin, 100 µg/mL streptomycin, and 2 mM L-glutamine (Gibco). Pooled human umbilical vein endothelial cells (HUVECs, Lonza, Basel, Switzerland) were cultured in endothelial cell growth medium 2 (EGM-2, Lonza)). For HUVEC passaging, tissue culture plates were precoated for 30 min at 37°C with 0.1% (w/v) porcine gelatin (Sigma, St. Louis, Missouri, US) in 1X PBS (Gibco). All cells were routinely tested for mycoplasma by PCR method.

### Transfection for Lentivirus Production

HEK-293T cells at 70-80% confluence were transfected using 3.3 µg of the target lentiviral construct, 2 µg pSPAX2 and 1 µg pMD2.G with 1 mg/mL polyethyenimine (PEI, Sigma) in serum-free DMEM. After 5 to 6 h, media was replaced with antibiotic-free DMEM containing 10% FBS and 2 mM L-glutamine (Gibco). Transfected cells were incubated for 48 h at 37°C to allow lentivirus production. The supernatant media containing viral particles was harvested and filtered through a 0.45 µm polyethersulfone (PES) filter (VWR, Randor, Pennsylvania, US) and aliquoted. Virus was stored at -80°C until use.

### Lentiviral Transduction

Lentivirus was supplied to wells of plated HUVECs in EGM-2 supplemented with 5 µg/mL hexadimethrine bromide (polybrene). After 24 h incubation, cells were selected with either 5 µg/mL blasticidin (Sigma) for 96 h, replacing the media and antibiotic at 48 h, or 1 µg/mL puromycin (Sigma) for 48 h. Following selection, HUVEC medium was replaced with EGM-2 without selection for at least 24 h recovery before further use. Lentivirus was titrated to ensure no significant changes in cell viability from non-targeting control.

### Immunofluorescence

Immunofluorescence was performed as described previously (Corcoran, Johnston, and McCormick 2015). Briefly, cells were grown on coverslips (no. 1.5, Electron Microscopy Sciences, Hatfield, Pennsylvania, US). Following treatment, coverslips were fixed in 4% paraformaldehyde (PFA, Electron Microscopy Sciences) in PBS at 37°C for 10 min, permeabilized with 0.1% Triton-X100 (Sigma) in 1X PBS for 10 min at RT, and blocked in 1% Human AB serum (blocking buffer, Sigma) in 1X PBS for 1 h at RT. Coverslips were then incubated with diluted primary antibody in blocking buffer overnight at 4°C in a humidified chamber. After primary antibody incubation, coverslips were washed with 1X PBS and then incubated in fluorescently-tagged secondary antibody diluted in blocking buffer for 1 h at RT. If applicable, coverslips were stained with Phalloidin-conjugated Alexa-Fluor 647 (Invitrogen, 1:100) in 1X PBS for 1.5 h. Coverslips were mounted onto microscope slides (FisherBrand, Pittsburgh, Pennsylvania, US) using Prolong Gold Antifade Mounting Media (Invitrogen, Carlsbad, California, US). For coverslips that were used for EDC4 puncta quantification, the following modifications to immunofluorescence were made: (1) Prior to permeabilization, coverslips were stained with wheat germ agglutinin (WGA) Alexa-647 conjugate (Invitrogen, 1:400) in 1X PBS for 10 min at RT. (2) Following secondary antibody incubation, coverslips were stained with 4’,6-Diamidino-2-Phenylindole (DAPI, Invitrogen, 1:10,000) in 1X PBS for 5 min.

Confocal imaging was performed on the Zeiss LSM 880 Confocal Microscope (Charbonneau Microscopy Facility, University of Calgary, Calgary, Canada) at the 63X oil objective. CellProfiler imaging was performed on the Zeiss AxioImager Z2 (CORES facility, Dalhousie University, Halifax, Canada) or Zeiss AxioObserver (Charbonneau Microscopy Facility, University of Calgary) at the 40X oil objective.

### Quantification of Processing Bodies Using CellProfiler Analysis

CellProfiler (cellprofiler.org) is an open source software for high-content image analysis (Kamentsky et al. 2011) and was used to develop an unbiased method for quantifying changes to PB dynamics. The pipeline used for quantifying PBs was structured as follows: To detect nuclei, the DAPI image was thresholded into a binary image. In the binary image, primary objects between 30 to 200 pixels in diameter were detected and defined as nuclei. Cells were identified as secondary objects in the WGA image using a propagation function from the identified nuclei, which determined the cell’s outer edge. Using the parameters of a defined nucleus and cell border, the cytoplasm was then defined as a tertiary object. The EDC4 channel image was enhanced using an “Enhance Speckles” function to identify distinct puncta and eliminate background staining. The cytoplasm image was then applied as a mask to the enhanced puncta image to ensure quantitation of only cytoplasmic puncta. EDC4 puncta were measured in the cytoplasm of cells using a ‘global thresholding with robust background adjustments’ function as defined by the program. The threshold cut-off for identified EDC4 puncta remained constant between all experiments with identical staining parameters. Puncta number per cell, intensity and locations with respect to the nucleus were measured and exported as .csv files and analyzed in RStudio. A template of the RStudio analysis pipeline is attached in Appendix A. Data was represented as fold change in EDC4 puncta count per cell normalized to the vector puncta count. ‘Relative EDC4 Puncta/Cell (KapB/Vector)’ demonstrates the KapB puncta count divided by vector puncta count, a ratio that was calculated within each treatment group for each biological replicate.

### Protein Electrophoresis and Immunoblotting

Cells were lysed in 2X Laemmli buffer (20% glycerol, 4% SDS, 120 mM Tris-HCl), between 150 to 300 µL, depending on cell density. Lysates were homogenized with a 0.21-gauge needle, and supplemented to contain 0.02% (w/v) bromophenol blue (Sigma) and 0.05 M dithiothreitol (DTT, Sigma), then heated at 95°C for 5 min. 7.5 or 12% TGX Stain-Free SDS-polyacrylamide gels (BioRad) were cast according to the instructions of the manufacturer and 5 to 15 µg of total protein were subjected to SDS gel electrophoresis using 1X SDS running buffer (25 mM Tris, 192 mM Glycine, 0.1% SDS). Precision Plus Protein All Blue Prestained Protein Standards (BioRad, Hercules, California, US) was used as a molecular weight marker. After electrophoresis, gels were UV-activated using the ChemiDocTouch (BioRad) Stain-Free Gel setting with automated exposure for 45 s. The protein was transferred to low-fluorescence polyvinylidene difluoride (PVDF) membranes (BioRad) on the Trans-Blot Turbo Transfer System (BioRad) according to the instructions of the manufacturer. Following transfer, total protein amounts on membranes were imaged on the ChemiDocTouch using the Stain-Free Membrane setting with automated exposure. Membranes were then blocked using 5% BSA (Sigma) in 1X TBS-T (150 nM NaCl, 10 mM Tris, pH 7.8, 0.01% Tween-20) for 1 h at RT. Primary antibody was diluted in 2.5% BSA in 1X TBS-T. Membranes were incubated in primary antibody solution overnight at 4°C with rocking. The following day, membranes were washed 3 times for 5 min in 1X TBS-T. Membranes were incubated with the appropriate secondary antibody, conjugated to horseradish peroxidase (HRP) for 1 h at RT. Membranes were washed 4 times for 5 min in 1X TBS-T. Clarity™ Western ECL Blotting Substrate (BioRad) was mixed at a 1:1 ratio and applied to the membrane for 5 min. Chemiluminescent signal was imaged on ChemiDocTouch Chemiluminescence setting. Band intensity was quantified using ImageLab software (BioRad), normalizing to total protein.

### Quantitative Reverse-Transcriptase Polymerase Chain Reaction (qRT-PCR)

Cells were lysed in 250 µL RLT buffer (Qiagen, Hilden, Germany) and RNA was extracted using the RNeasy Plus Mini kit (Qiagen) according to the manufacturer’s instructions. Complementary DNA (cDNA) was synthesized from 1 µg of total RNA using the qScript cDNA SuperMix (QuantaBio, Hilden, Germany) according to the manufacturer’s instructions. Real-time quantitative PCR with SsoFast EvaGreen qPCR MasterMix (BioRad) was used to quantify the fold-change in mRNA abundance. Relative fluorescence was quantified using CFX Connect (BioRad). All qRT-PCR primers efficiencies were between 90-110% in HUVECs and sequences are found in Table S5.

### Collagen-Coating for Altering Matrix Stiffness

Coverslips (no. 1.5, Electron Microscopy Sciences) were coated with a dilution series (0 to 64 µg/cm^2^) of rat-tail collagen-1 (Gibco) in 0.02 M acetic acid for 4 h at RT. Slides were sterilized with UV irradiation and washed with 2 times with sterile 1X PBS prior to seeding cells.

### Unidirectional Fluid Flow for Endothelial Cell Shear Stress

A parallel-plate flow chamber was used to expose ECs to shear stress. The system was described in detail in Gomez-Garcia et al. (2018). Briefly, cleaned, unfrosted microscope slides (Cole-Parmer, Vernon Hills, Illinois, US) were coated for 4 h at RT with rat-tail collagen-1 (Gibco) in 0.02 M acetic acid for a resultant 8.3 µg/cm^2^ collagen density. Slides were sterilized with UV irradiation and washed 2 times with sterile 1X PBS. HUVECs were seeded at a density of 350,000 cells/slide and cultured for 24 h. Forty-five mL of EGM-2 supplemented with dextran (Spectrum Chemical, New Brunswick, New Jersey, US) for a resultant 3 cP viscosity was added to the stock media bottle. The stock media bottle was connected with the associated tubing and pulse dampener. Slides with seeded cells were inserted onto the flow chamber, a gasket (Specialty Manufacturing, Calgary, Canada) was added, and the system was sealed shut and attached to the flow loop following the outlet of a pulse dampener. The rate of fluid flow was started at 0.3 L/min and doubled every 15 min until final flow rates of 0.6 L/min and 2.7 L/min were reached, corresponding to shear stress rates of 2 and 10 dyn/cm^2^. Following 21 h, cells were removed and immediately fixed for immunofluorescence or lysed for immunoblotting.

### Statistical Analysis

Statistical analysis was performed in GraphPad Prism 8.0 software. Significance was determined using a ratio paired t-test or repeated measures analysis of variance (ANOVA). One-tailed ratio paired t-tests were applied in comparisons specifically examining PB restoration in KapB-expressing cells as a directional hypothesis. In all other comparisons, two-tailed ratio paired t-tests were applied. Significance was determined at *p* = 0.05. Each biological replicate for CellProfiler quantification consisted of 6 images of each treatment in a given experiment, counting approximately 100 to 200 cells per treatment.

## Acknowledgements

We would like to sincerely thank the members of the Corcoran lab for helpful discussions about this work. We would like to thank Dr. Craig McCormick (Dalhousie University) for helpful discussions on this manuscript and his lab for plasmids, expertise and invaluable advice. We would like to thank Mr. Stephen Whitefield of the Dalhousie *CORES* Cellular and Molecular Digital Imaging Facility and Dr. Anne Vaahtokari of the Charbonneau Microscopy Facility, UCalgary for microscopy support. ELC was supported by a Killam predoctoral scholarship, an NSERC CGS-M scholarship, and a Nova Scotia Graduate scholarship. Operating funds to support this work derive from an NSERC Discovery grant RGPIN-2015-04882 to JAC.

## Supplementary Files

Table 1: Table detailing antibodies used in this study, including species, target, source, use and dilution.

Table 2: Table detailing plasmids used in this study, including name, use, source, bacterial selection cassette and mammalian selection cassette.

Table 3: Table detailing shRNA sequences used in this study.

Table 4: Table detailing drug treatments used in this study, including name, use, source, concentration and duration of treatment.

Table 5: Table detailing qRT-PCR primers used in this study, including target, sequence, melting temperature, and reference.

## Source Code

**Source Code 1**: CellProfiler pipeline used to enumerate puncta from epifluorescent microscopy images of HUVECs stained for Hedls puncta; co-stained with DAPI and WGA.

**Source Code 2**: CellProfiler pipeline used to enumerate puncta from confocal microscopy images of HUVECs subjected to shear stress stained for Hedls and DDX6 puncta.

**Source Code 3**: CellProfiler pipeline used to enumerate puncta from confocal microscopy images of HUVECs subjected to shear stress stained for Hedls puncta.

**Source Code 4**: CellProfiler pipeline used to quantify intensity in nucleus and cytoplasm from epifluorescent microscopy images of HUVECs stained for YAP protein; co-stained with DAPI and WGA.

**Source Code 5:** Template RStudio code used for analysis of CellProfiler output from Source Code 1. Annotations provided within code.

**Source Code 6:** Template RStudio code used for analysis of CellProfiler output from Source Code 2. Annotations provided within code.

**Source Code 7:** Template RStudio code used for analysis of CellProfiler output from Source Code 3. Annotations provided within code.

**Source Code 8:** Template RStudio code used for analysis of CellProfiler output from Source Code 4. Annotations provided within code.

---

**Figure S1:**
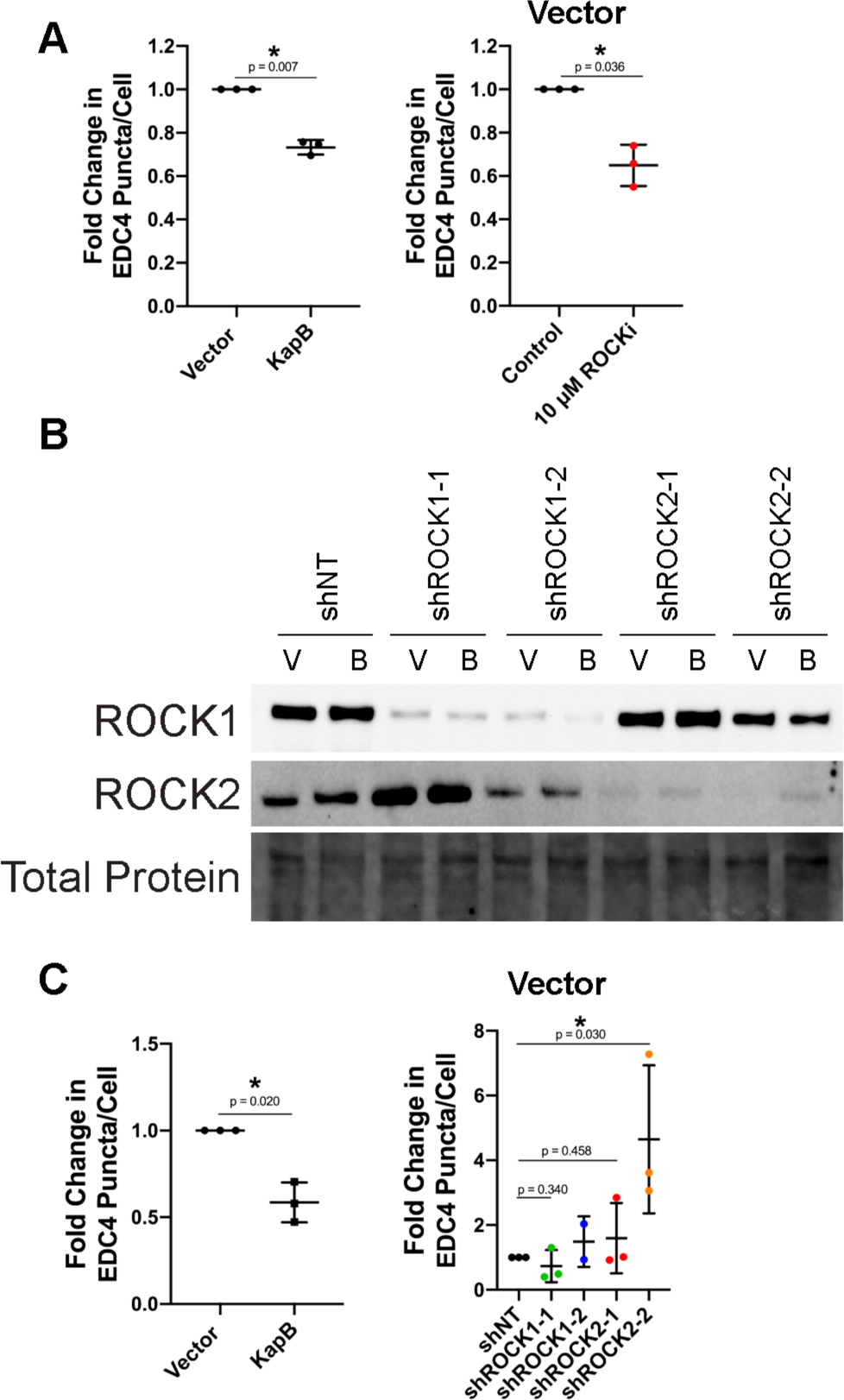
The RhoA-effector ROCK is required for KapB-mediated PB disassembly, knockdown confirmation and vector data. (A) KapB- and vector- expressing HUVECs were treated with 10 µM Y-27632 or water control for 4 h and fixed for immunofluorescence. Fixed cells were stained for CellProfiler analysis as detailed in the methods. The number of EDC4 puncta per cell was quantified and normalized to the vector control. Vector control data is shown. (B, C) KapB- and vector- expressing HUVECs were transduced with shRNAs targeting ROCK1 and ROCK2 (shROCK1-1, shROCK1-2, shROCK2-1, shROCK2-2) or with a non- targeting (shNT) control and selected. In parallel, cells were lysed for immunoblotting or fixed for immunofluorescence. (B) One representative immunoblot of three independent experiments stained using ROCK1- and 2-specific antibodies. (C) Fixed cells were stained for CellProfiler analysis as detailed in the methods. The number of EDC4 puncta per cell was quantified and normalized to the vector NT control within each replicate. Vector control data is shown. Statistics were determined using a ratio paired t-test between control and experimental groups; error bars represent standard deviation; n=3 independent biological replicates; * = p < 0.05.

**Figure S2:**
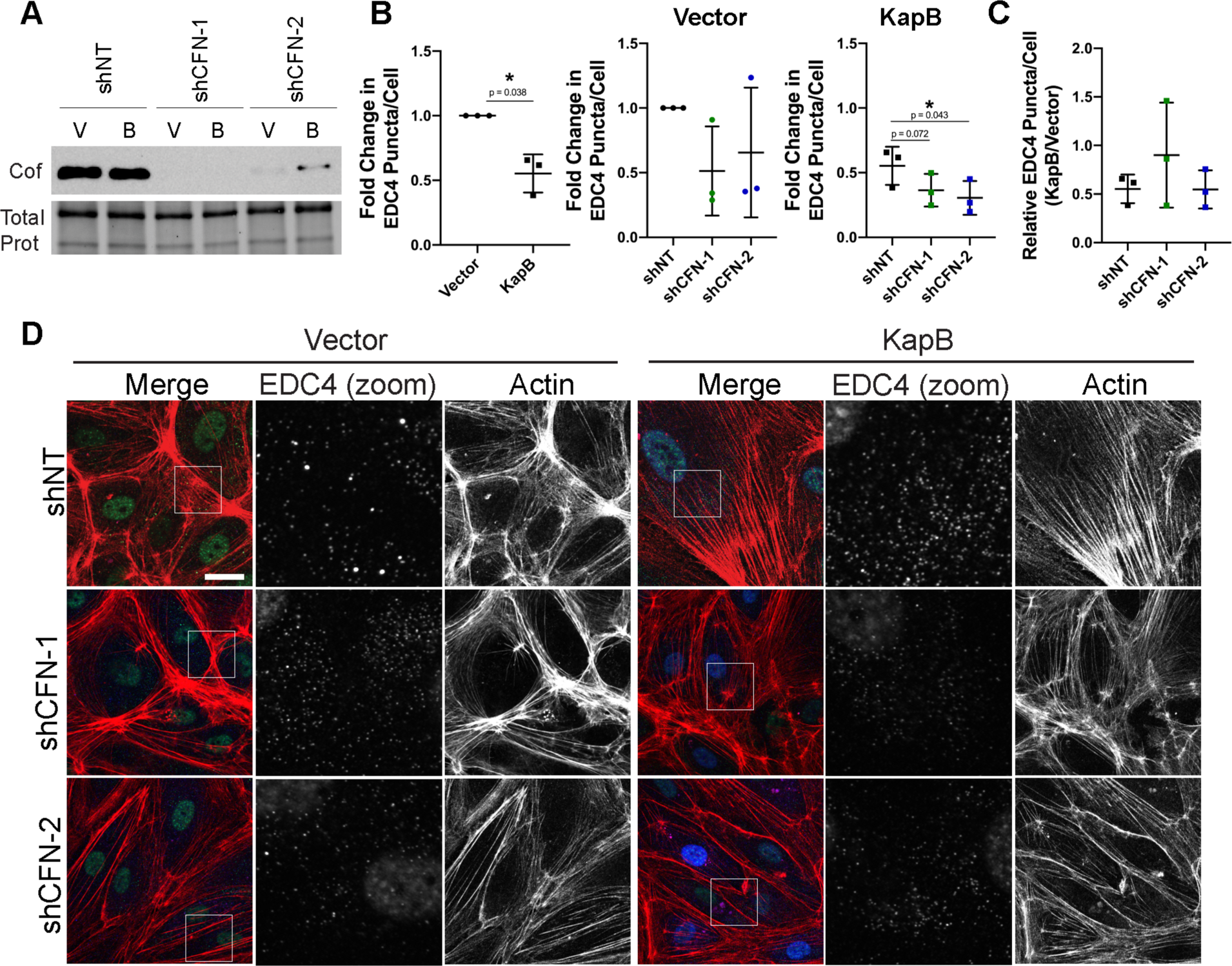
Cofilin knockdown augments KapB-mediated PB disassembly. KapB- and vector- expressing HUVECs were transduced with shRNAs targeting cofilin (shCFN-1, shCFN- 2) or with a non-targeting (shNT) control and selected. In parallel, cells were fixed for immunofluorescence or lysed for immunoblotting. (A) One representative immunoblot of three independent experiments stained using a cofilin-specific antibody. (B, C) Fixed cells were stained for CellProfiler analysis as detailed in the methods. (B) The number of EDC4 puncta per cell was quantified and normalized to the vector NT control within each replicate. (C) CellProfiler data was used to calculate the ratio of EDC4 puncta counts in KapB-expressing cells versus the vector control for each treatment condition. (D) Representative images of cells stained for PB-resident protein EDC4 (green), KapB (blue), and F-actin (red, phalloidin). Boxes indicate the area of the field of view that is shown in EDC4 (zoom) panel. Scale bar represents 20 µm. Statistics were determined using a ratio paired t-test between control and experimental groups; error bars represent standard deviation; n=3 independent biological replicates; * = p < 0.05.

**Figure S3:**
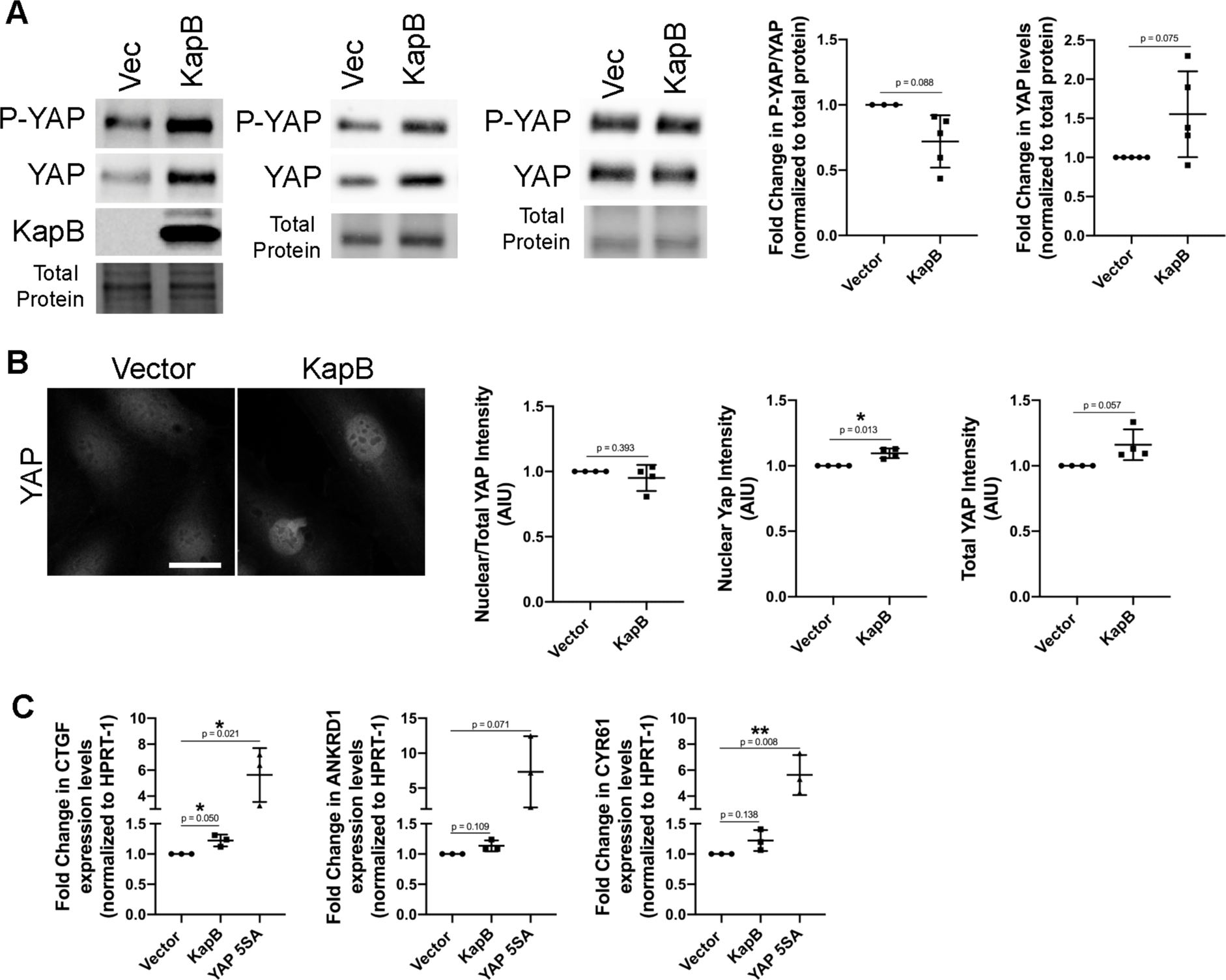
KapB does not activate canonical functions of YAP. (A, B) KapB- and vector- expressing HUVECs were lysed for immunoblotting or fixed for immunofluorescence. (A) Representative immunoblots and quantification of immunoblotting for P(S127)-YAP-, YAP- or KapB-specific antibody are shown. Several immunoblots are shown to illustrate variation in KapB-mediated changes in P-YAP and YAP. Protein levels in each condition were normalized to total protein. All treatments were normalized to vector control within each replicate. (B) Representative images of cells stained for YAP. Scale bar represents 20 µM. (C) HUVECs were transduced with recombinant lentiviruses expressing KapB, a constitutively active version of YAP (YAP 5SA) or an empty vector control, selected and lysed for total RNA. qRT-PCR analysis of steady state mRNA levels of canonical YAP-regulated genes CTGF, ANKRD1 and CYR61 was performed, and was normalized to steady state HPRT-1 mRNA levels. Statistics were determined using repeated measures ANOVA; error bars represent standard deviation; n=3 independent biological replicates; * = p < 0.05, ** = p < 0.01.

**Supplementary Table 1:** Antibodies used in this study.

| **Antibody** | **Source** | **Use** | **Dilution** |
| --- | --- | --- | --- |
| Rabbit α-mDia1 | Cell Signalling Technologies<br>(Cat#:5486) | Immunoblotting | 1:1000 in 2.5% BSA |
| Rabbit α-ROCK1 | Cell Signalling Technologies<br>(Cat#:4035) | Immunoblotting | 1:1000 in 2.5% BSA |
| Rabbit α-ROCK2 | Cell Signalling Technologies<br>(Cat#:9029) | Immunoblotting | 1:500 in 2.5% BSA |
| Rabbit α-Cofilin | Cell Signalling Technologies<br>(Cat#:5175) | Immunoblotting | 1:1000 in 2.5% BSA |
| Rabbit α-P-YAP | Cell Signalling Technologies<br>(Cat#: 4911) | Immunoblotting | 1:1000 in 2.5% BSA |
| Rabbit α-YAP | Cell Signalling Technologies<br>(Cat#: 4912) | Immunoblotting | 1:1000 in 2.5% BSA |
| α-Mouse IgG, HRP-linked (2°) | Cell Signalling Technologies<br>(Cat#: 7076) | Immunoblotting | 1:2000 to 1:4000 in 2.5% BSA |
| α-Rabbit IgG, HRP-linked (2°) | Cell Signalling Technologies<br>(Cat#: 7074) | Immunoblotting | 1:2000 to 1:4000 in 2.5% BSA |
| Mouse α-p70 s6 kinase (detects EDC4) | Santa Cruz (Cat#:sc-8418) | Immunofluorescence | 1:1000 in blocking buffer (1% Human AB in PBS), 4°C overnight |
| Rabbit α-KapB | Gift from D. Ganem and C. McCormick | Immunofluorescence/ Immunoblotting | 1:1000 in blocking buffer (1% Human AB in PBS), 30 min RT |
| Mouse α-YAP | Santa Cruz (Cat#:sc-101199) | Immunofluorescence | 1:1000 in blocking buffer (1% Human AB in PBS), 4°C overnight |
| Rabbit α-DDX6 | Bethyl Labs (Cat#:A300-461A) | Immunofluorescence | 1:1000 in blocking buffer (1% Human AB in PBS), 4°C overnight |
| Rabbit α-β-actin HRP-linked | Cell Signalling Technologies (Cat #:5125) | Immunoblotting | 1:2000 in 2.5% BSA |
| Alexa Fluor 555-conjugated donkey α-mouse IgG (2°) | Invitrogen (Cat#:A31570) | Immunofluorescence | 1:1000 in blocking buffer (1% Human AB in PBS), 1h RT |
| Alexa Fluor 488-conjugated chicken α-rabbit IgG (2°) | Invitrogen (Cat#:A21441) | Immunofluorescence | 1:1000 in blocking buffer (1% Human AB in PBS), 1h RT |
| Alexa Fluor 555-conjugated donkey α-rabbit IgG (2°) | Invitrogen (Cat#:A31572) | Immunofluorescence | 1:1000 in blocking buffer (1% Human AB in PBS), 1h RT |
| Alexa Fluor 488-conjugated chicken α-mouse IgG (2°) | Invitrogen (Cat#:A21200) | Immunofluorescence | 1:1000 in blocking buffer (1% Human AB in PBS), 1h RT |

**Supplementary Table 2:** Plasmids used in this study.

| **Plasmid Name** | **Use** | **Source** | **Bacterial Selection Cassette** | **Mammalian Selection Cassette (Lentiviral Plasmids Only)** |
| --- | --- | --- | --- | --- |
| pLJM-1-EV | Control vector for lentiviral expression studies | C. McCormick (Dalhousie University) | Ampicillin | Blasticidin, Puromycin |
| pLJM-1 KapB | Lentiviral expression of KapB | C. McCormick (Dalhousie University) | Ampicillin | Blasticidin |
| pLKO-(shRNA) | Lentiviral expression of short hairpin RNAs (shRNA sequences in Table S3) | Cloned from: pLKO-TRC Addgene no.: 26655 | Ampicillin | Puromycin |
| pLJM-1-YAP-5SA (CA-YAP) | Lentiviral expression of constitutively active YAP | Cloned from: p2XFLAG-YAP-5SA, Donated by C. McCormick (Dalhousie University) | Ampicillin | Blasticidin |
| pcDNA3.1 | Transfection control | Invitrogen | Ampicillin | N/A |
| pcDNA3.1 KapB | Transfection of KapB | C. McCormick (Dalhousie University) | Ampicillin | N/A |
| p1XFLAG | Transfection control | Cloned from: p2XFLAG-YAP-5SA, Donated by C. McCormick (Dalhousie University) | Ampicillin | N/A |
| p2XFLAG-YAP 5SA | Transfection of YAP 5SA | Donated by C. McCormick (Dalhousie University) | Ampicillin | N/A |
| pV5-YAP 6SA | Transfection of YAP 6SA | Addgene no: 42562 | Ampicillin | N/A |
| pMD2.G | Envelope protein for lentiviral production | Addgene no: 12259 | Ampicillin | N/A |
| psPAX2 | Packaging proteins for lentiviral production | Addgene no: 12260 | Ampicillin | N/A |

**Supplementary Table 3:**
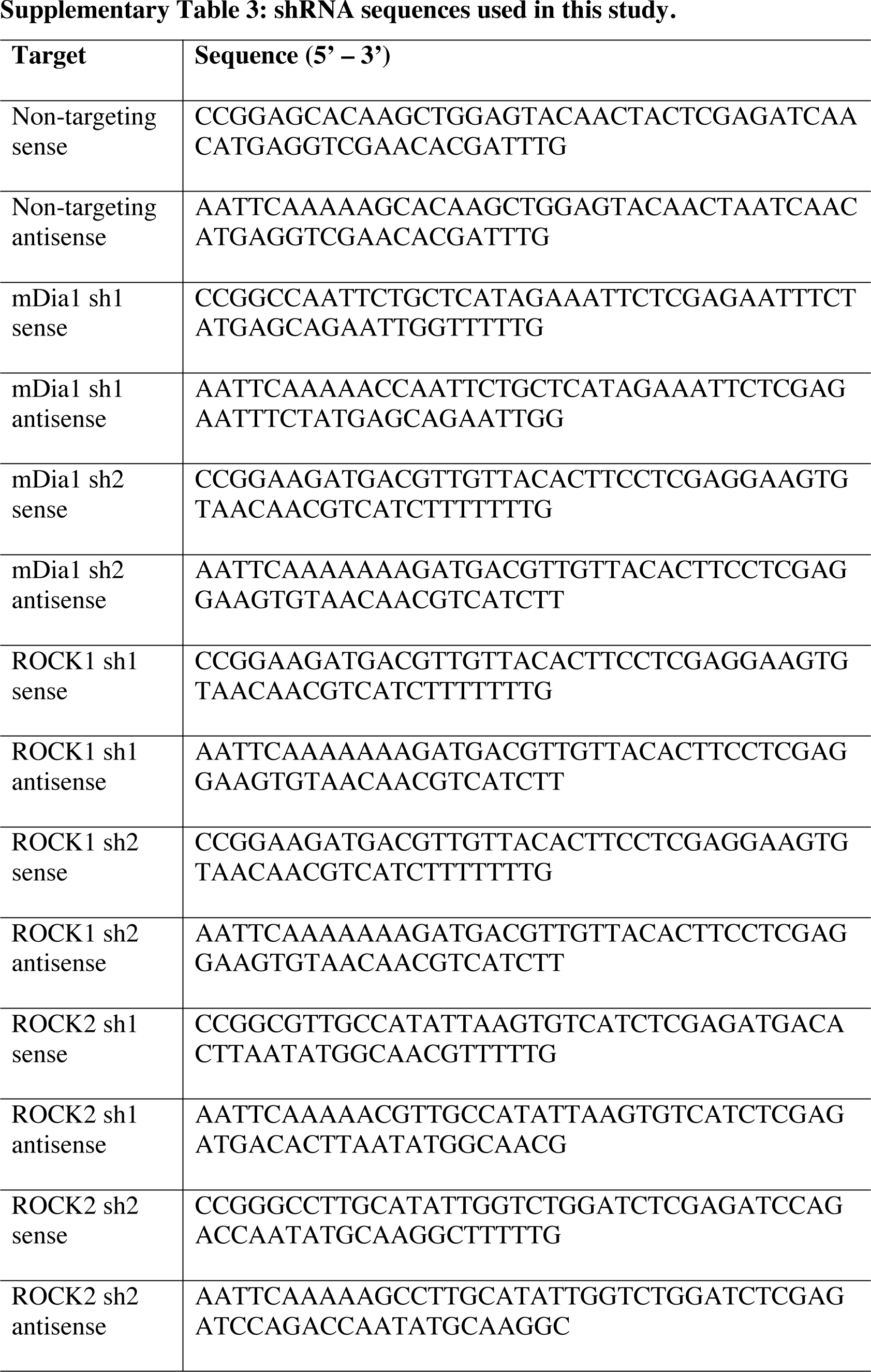
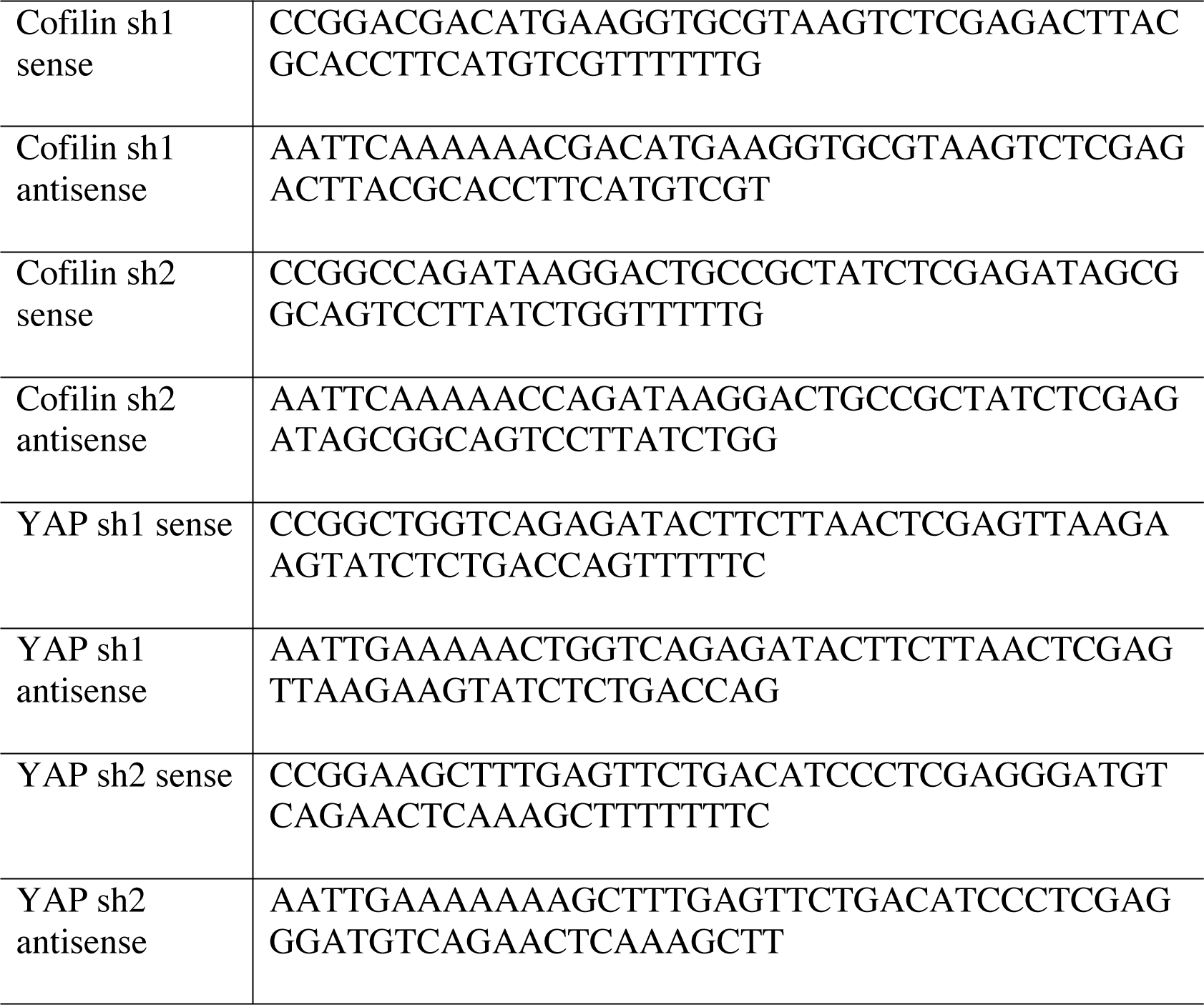
shRNA sequences used in this study.

**Supplementary Table 4:** Drug treatments used in this study.

| **Drug** | **Use** | **Source (Cat#)** | **Concentration Used** | **Duration** |
| --- | --- | --- | --- | --- |
| Y-27623 dihydrochloride (ROCKi) | Non-isoform specific inhibition of ROCK | Sigma-Aldrich (Cat#:Y0503) | 10 μM | 4 h |
| (-)-Blebbistatin | Inhibition of MLC contractility | Sigma-Aldrich (Cat#:B0560) | 10 μM | 30 min |
| Calyculin A | Inhibition of MLC phosphatase, resulting in cell contraction | Abcam (Cat#: ab141784) | 2.5 nM, 5 nM | 20 min |

**Supplementary Table 5:**
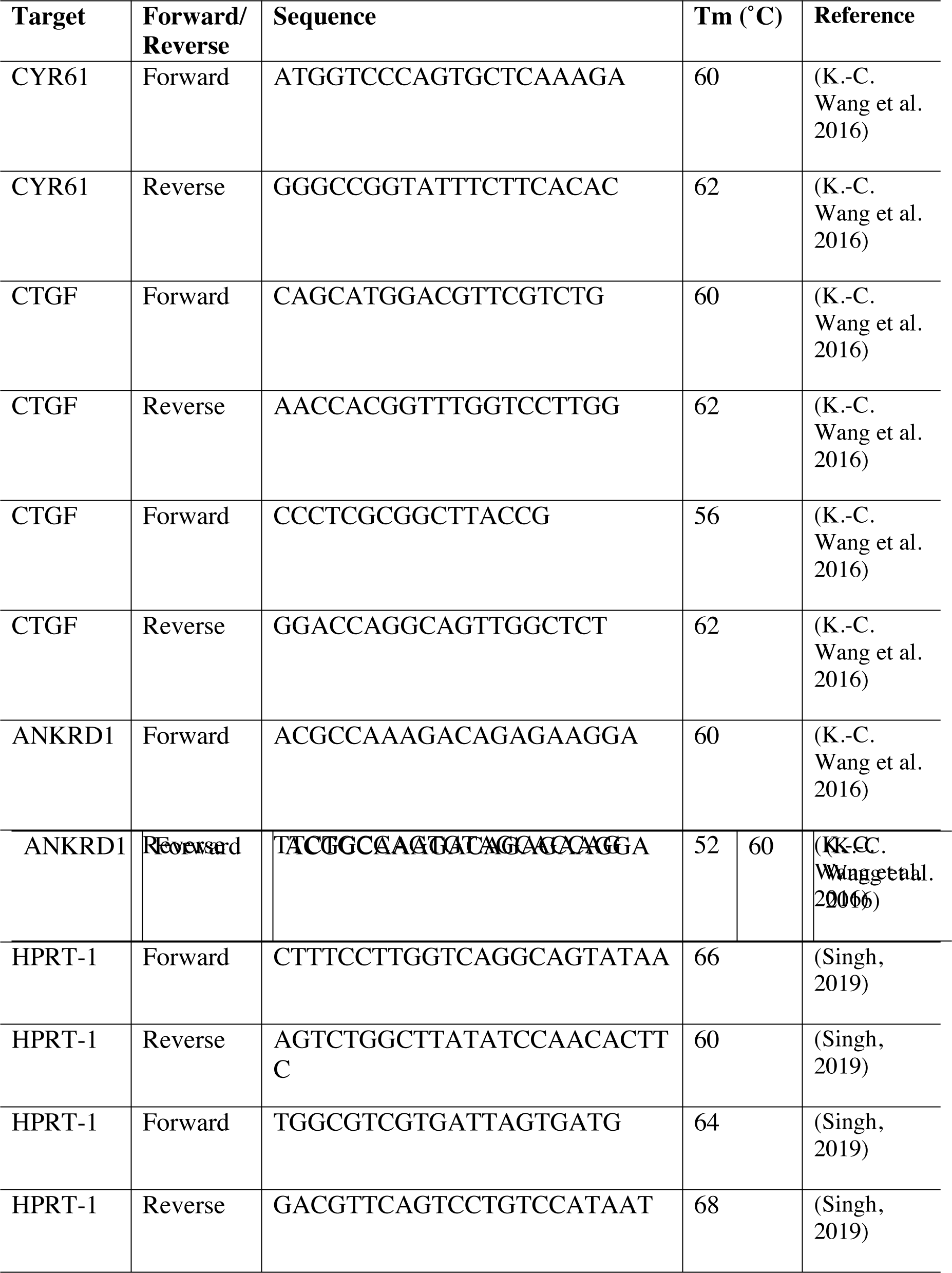
qRT-PCR primers used in this study.

## References

1. Aizer, Adva, Yehuda Brody, Lian Wee Ler, Nahum Sonenberg, Robert H. Singer, and Yaron Shav-Tal. 2008. “The Dynamics of Mammalian P Body Transport, Assembly, and Disassembly In Vivo.” Molecular Biology of the Cell 19 (11): 4154–66. https://doi.org/10.1091/mbc.E08.

2. Amano, M, K Chihara, K Kimura, Y Fukata, N Nakamura, Y Matsuura, and K Kaibuchi. 1997. “Formation of Actin Stress Fibers and Focal Adhesions Enhanced by Rho-Kinase.” Science 275 (5304): 1308–11. https://doi.org/10.1126/science.275.5304.1308.

3. Amano, Mutsuki, Masaaki Ito, Kazushi Kimura, Yuko Fukata, Kazuyasu Chihara, Takeshi Nakano, Yoshiharu Matsuura, and Kozo Kaibuchi. 1996. “Phosphorylation and Activation of Myosin by Rho-Associated Kinase (Rho-Kinase).” The Journal of Biological Chemistry 271 (34): 20246–49. https://doi.org/10.1074/jbc.271.34.20246.

4. Arias, Carolina, Ben Weisburd, Noam Stern-Ginossar, Alexandre Mercier, Alexis S Madrid, Priya Bellare, Meghan Holdorf, Jonathan S. Weissman, and Don Ganem. 2014. “KSHV 2.0: A Comprehensive Annotation of the Kaposi’s Sarcoma-Associated Herpesvirus Genome Using Next-Generation Sequencing Reveals Novel Genomic and Functional Features.” PLoS Pathogens 10 (1). https://doi.org/10.1371/journal.ppat.1003847.

5. Asano, Yukako, and Issei Mabuchi. 2001. “Calyculin-A, an Inhibitor for Protein Phosphatases, Induces Cortical Contraction in Unfertilized Sea Urchin Eggs.” Cell Motility and the Cytoskeleton 48 (4): 245–61. https://doi.org/10.1002/cm.1013.

6. Bakheet, Tala, Edward Hitti, and Khalid S A Khabar. 2017. “ARED-Plus: An Updated and Expanded Database of AU-Rich Element-Containing MRNAs and Pre-MRNAs.” Nucleic Acids Research 46 (October 2017): 2017–19. https://doi.org/10.1093/nar/gkx975.

7. Bakheet, Tala, Bryan R G Williams, and Khalid S A Khabar. 2006. “ARED 3.0: The Large and Diverse AU-Rich Transcriptome.” Nucleic Acids Research 34 (Database issue): D111–14. https://doi.org/10.1093/nar/gkj052.

8. Balinsky, Corey A, Hana Schmeisser, Alexandra I Wells, Sundar Ganesan, Tengchuan Jin, Kavita Singh, and Kathryn C Zoon. 2017. “IRAV (FLJ11286), an Interferon-Stimulated Gene with Antiviral Activity against Dengue Virus, Interacts with MOV10.” Edited by Michael S. Diamond. Journal of Virology 91 (5). https://doi.org/10.1128/JVI.01606-16.

9. Blanco, Fernando F, Sandhya Sanduja, Natasha G Deane, Perry J Blackshear, and Dan A Dixon. 2014. “Transforming Growth Factor β Regulates P-Body Formation through Induction of the MRNA Decay Factor Tristetraprolin.” Molecular and Cellular Biology 34 (2): 180–95. https://doi.org/10.1128/MCB.01020-13.

10. Boshoff, Chris, Thomas F Schulz, Margaret M Kennedy, Andrew K Graham, Cyril Fisher, Alero Thomas, J O’D. McGee, Robin a Weiss, and John J O’Leary. 1995. “Kaposi’s Sarcoma-Associated Herpesvirus Infects Endothelial and Spindle Cells.” Nature Medicine 1 (12): 1274–78. https://doi.org/10.1038/nm1295-1274.

11. Burdick, Ryan, Jessica L Smith, Chawaree Chaipan, Yeshitila Friew, Jianbo Chen, Narasimhan J Venkatachari, Krista A Delviks-Frankenberry, Wei-Shau Hu, and Vinay K. Pathak. 2010. “P Body-Associated Protein Mov10 Inhibits HIV-1 Replication at Multiple Stages.” Journal of Virology 84 (19): 10241–53. https://doi.org/10.1128/jvi.00585-10.

12. Burgess, Hannah M., and Ian Mohr. 2015. “Cellular 5′-3′ MRNA Exonuclease Xrn1 Controls Double-Stranded RNA Accumulation and Anti-Viral Responses.” Cell Host and Microbe 17 (3): 332–44. https://doi.org/10.1016/j.chom.2015.02.003.

13. Burridge, Keith, and Christophe Guilluy. 2016. “Focal Adhesions, Stress Fibers and Mechanical Tension.” Experimental Cell Research 343 (1): 14–20. https://doi.org/10.1016/j.yexcr.2015.10.029.

14. Calvo, Fernando, Nil Ege, Araceli Grande-Garcia, Steven Hooper, Robert P. Jenkins, Shahid I. Chaudhry, Kevin Harrington, et al. 2013. “Mechanotransduction and YAP-Dependent Matrix Remodelling Is Required for the Generation and Maintenance of Cancer-Associated Fibroblasts.” Nature Cell Biology 15 (6): 637–46. https://doi.org/10.1038/ncb2756.

15. Chang, Y, E Cesarman, M S Pessin, F Lee, J Culpepper, D M Knowles, and P S Moore. 1994. “Identification of Herpes-like DNA Sequences in AIDS-Associated Kaposi’s Sarcoma.” Science 266: 1865–69. https://doi.org/10.1126/science.7997879.

16. Chen, C Y, and A B Shyu. 1995. “AU-Rich Elements: Characterization and Importance in MRNA Degradation.” Trends in Biochemical Sciences 20 (11): 465–70. https://doi.org/10.1016/S0968-0004(00)89102-1.

17. Ciufo, D M, J S Cannon, L J Poole, F Y Wu, P Murray, R F Ambinder, and G S Hayward. 2001. “Spindle Cell Conversion by Kaposi’s Sarcoma-Associated Herpesvirus: Formation of Colonies and Plaques with Mixed Lytic and Latent Gene Expression in Infected Primary Dermal Microvascular EndCiufo, D. M., Cannon, J. S., Poole, L. J., Wu, F. Y., Murray, P.,.” Journal of Virology 75 (12): 5614–26. https://doi.org/10.1128/JVI.75.12.5614-5626.2001.

18. Corcoran, Jennifer A, Benjamin P Johnston, and Craig McCormick. 2015. “Viral Activation of MK2-Hsp27-P115RhoGEF-RhoA Signaling Axis Causes Cytoskeletal Rearrangements, P-Body Disruption and ARE-MRNA Stabilization.” Edited by Erle S. Robertson. PLoS Pathogens 11 (1): e1004597. https://doi.org/10.1371/journal.ppat.1004597.

19. Corcoran, Jennifer A, Denys A Khaperskyy, Benjamin P Johnston, Christine A King, David P. Cyr, Alisha V. Olsthoorn, and Craig McCormick. 2012. “Kaposi’s Sarcoma-Associated Herpesvirus G-Protein-Coupled Receptor Prevents AU-Rich-Element-Mediated MRNA Decay.” Journal of Virology 86 (16): 8859–71. https://doi.org/10.1128/JVI.00597-12.

20. Corcoran, Jennifer A, and Craig McCormick. 2015. “Viral Activation of Stress-Regulated Rho-GTPase Signaling Pathway Disrupts Sites of MRNA Degradation to Influence Cellular Gene Expression.” Small GTPases 6 (4): 178–85. https://doi.org/10.1080/21541248.2015.1093068.

21. Cramer, Louise P, Margaret Siebert, and Timothy J Mitchison. 1997. “Identification of Novel Graded Polarity Actin Filament Bundles in Locomoting Heart Fibroblasts: Implications for the Generation of Motile Force.” Journal of Cell Biology 136 (6): 1287–1305. https://doi.org/10.1083/jcb.136.6.1287.

22. Cuevas, Rolando A, Arundhati Ghosh, Christina Wallerath, Veit Hornung, Carolyn B Coyne, and Saumendra N Sarkar. 2016. “MOV10 Provides Antiviral Activity against RNA Viruses by Enhancing RIG-I–MAVS-Independent IFN Induction.” The Journal of Immunology 196 (9): 3877–86. https://doi.org/10.4049/jimmunol.1501359.

23. DiMaio, TA, KD Gutierrez, M. Lagunoff. 2011. Latent KSHV infection of endothelial cells induces integrin beta3 to activate angiogenic phenotypes. PLoS Pathog. 7:e1002424.

24. Docena, G, L Rovedatti, L Kruidenier, Á Fanning, NAB Leakey, CH Knowles, K Lee, et al. 2010. “Down-Regulation of P38 Mitogen-Activated Protein Kinase Activation and Proinflammatory Cytokine Production by Mitogen-Activated Protein Kinase Inhibitors in Inflammatory Bowel Disease.” Clinical and Experimental Immunology 162 (1): 108–15. https://doi.org/10.1111/j.1365-2249.2010.04203.x.

25. Dupont, Sirio, Leonardo Morsut, Mariaceleste Aragona, Elena Enzo, Stefano Giulitti, Michelangelo Cordenonsi, Francesca Zanconato, et al. 2011. “Role of YAP/TAZ in Mechanotransduction.” Nature 474 (7350): 179–83. https://doi.org/10.1038/nature10137.

26. Ensoli, B. 1998. “Kaposi’s Sarcoma: A Result of the Interplay among Inflammatory Cytokines, Angiogenic Factors and Viral Agents.” Cytokine & Growth Factor Reviews 9 (1): 63–83. https://doi.org/10.1016/S1359-6101(97)00037-3.

27. Eulalio, Ana, Isabelle Behm-Ansmant, and Elisa Izaurralde. 2007. “P Bodies: At the Crossroads of Post-Transcriptional Pathways.” Nature Reviews Molecular Cell Biology 8 (1): 9–22. https://doi.org/10.1038/nrm2080.

28. Finch-Edmondson, Megan, and Marius Sudol. 2016. “Framework to Function: Mechanosensitive Regulators of Gene Transcription.” Cellular & Molecular Biology Letters 21 (1): 28. https://doi.org/10.1186/s11658-016-0028-7.

29. Franks, Tobias M, and Jens Lykke-Andersen. 2007. “TTP and BRF Proteins Nucleate Processing Body Formation to Silence MRNAs with AU-Rich Elements.” Genes and Development 21 (6): 719–35. https://doi.org/10.1101/gad.1494707.

30. Friedland, Julie C, Mark H Lee, and David Boettiger. 2009. “Mechanically Activated Integrin Switch Controls Alpha5 Beta1 Function.” Science 323 (January): 642–44.

31. “G-Actin / F-Actin In Vivo Assay Kit.” n.d. Cytoskeleton Inc.

32. Ganem, Don. 1997. “KSHV and Kaposi’s Sarcoma: The End of the Beginning?” Cell 91 (2): 157–60. https://doi.org/10.1016/S0092-8674(00)80398-0.

33. Garcia, Melissa C, Denise M Ray, Brad Lackford, Mark Rubino, Kenneth Olden, and John D Roberts. 2009. “Arachidonic Acid Stimulates Cell Adhesion through a Novel P38 MAPK- RhoA Signaling Pathway That Involves Heat Shock Protein 27.” Journal of Biological Chemistry 284 (31): 20936–45. https://doi.org/10.1074/jbc.M109.020271.

34. Gill, MB, JE Murphy, JD Fingeroth. 2005. Functional divergence of Kaposi’s sarcoma-associated herpesvirus and related gamma-2 herpesvirus thymidine kinases: novel cytoplasmic phosphorylation that alter cellular morphology and disrupt adhesion. J Virol. 79:14647–59.

35. Gill, MB, R. Turner, PG Stevenson, M. Way. 2015. KSHV-TK is a tyrosine kinase that disrupts focal adhesions and induces Rho-mediated cell contraction. EMBO J. 34:448–465.

36. Gomez-Garcia, M Juliana, Amber L Doiron, Robyn RM Steele, Hagar I Labouta, Bahareh Vafadar, Robert D Shepherd, Ian D Gates, David T Cramb, Sarah J Childs, and Kristina D Rinker. 2018. “Nanoparticle Localization in Blood Vessels: Dependence on Fluid Shear Stress, Flow Disturbances, and Flow-Induced Changes in Endothelial Physiology.” Nanoscale 10 (32): 15249–61. https://doi.org/10.1039/C8NR03440K.

37. Grashoff, Carsten, Brenton D Hoffman, Michael D Brenner, Ruobo Zhou, Maddy Parsons, Michael T Yang, Mark A McLean, et al. 2010. “Measuring Mechanical Tension across Vinculin Reveals Regulation of Focal Adhesion Dynamics.” Nature 466 (7303): 263–66. https://doi.org/10.1038/nature09198.

38. Grossmann, Claudia, Simona Podgrabinska, Mihaela Skobe, and Don Ganem. 2006. “Activation of NF-ΚB by the Latent VFLIP Gene of Kaposi’s Sarcoma-Associated Herpesvirus Is Required for the Spindle Shape of Virus-Infected Endothelial Cells and Contributes to Their Proinflammatory Phenotype.” Journal of Virology 80 (14): 7179–85. https://doi.org/10.1128/jvi.01603-05.

39. Halder, Georg, Sirio Dupont, and Stefano Piccolo. 2012. “Transduction of Mechanical and Cytoskeletal Cues by YAP and TAZ.” Nature Publishing Group 13 (9): 591–600. https://doi.org/10.1038/nrm3416.

40. Horvathova I, Voigt F, Kotrys A V., Zhan Y, Artus-Revel CG, Eglinger J, et al. 2017. “The Dynamics of mRNA Turnover Revealed by Single-Molecule Imaging in Single Cells.” Molecular Cell 68 (3):615–625. https://doi.org/10.1016/j.molcel.2017.09.030

41. Hotulainen, Pirta, and Pekka Lappalainen. 2006. “Stress Fibers Are Generated by Two Distinct Actin Assembly Mechanisms in Motile Cells.” Journal of Cell Biology 173 (3): 383–94. https://doi.org/10.1083/jcb.200511093.

42. Huang, Yu, Li Wang, Jiang Yun Luo, Bochuan Li, Xiao Yu Tian, Li Jing Chen, Yuhong Huang, et al. 2016. “Integrin-YAP/TAZ-JNK Cascade Mediates Atheroprotective Effect of Unidirectional Shear Flow.” Nature 540 (7634): 579–82. https://doi.org/10.1038/nature20602.

43. Hubstenberger, Arnaud, Maïté Courel, Marianne Bénard, Sylvie Souquere, Michèle Ernoult- Lange, Racha Chouaib, Zhou Yi, et al. 2017. “P-Body Purification Reveals the Condensation of Repressed MRNA Regulons.” Molecular Cell 68 (1): 144–157.e5. https://doi.org/10.1016/j.molcel.2017.09.003.

44. Ishizaki, T, M Uehata, I Tamechika, J Keel, K Nonomura, M Maekawa, and S Narumiya. 2000. “Pharmacological Properties of Y-27632, a Specific Inhibitor of Rho-Associated Kinases.” Molecular Pharmacology 57 (5): 976–83. http://www.ncbi.nlm.nih.gov/pubmed/10779382.

45. Jackson, Wesley M, Michael J Jaasma, Andrew D Baik, and Tony M Keaveny. 2008. “Over- Expression of Alpha-Actinin with a GFP Fusion Protein Is Sufficient to Increase Whole- Cell Stiffness in Human Osteoblasts.” Annals of Biomedical Engineering 36 (10): 1605–14. https://doi.org/10.1007/s10439-008-9533-9.

46. Jang, Wonyul, Tackhoon Kim, Ja Seung Koo, Sang-kyum Kim, and Dae-Sik Lim. 2017. “ Mechanical Cue-induced YAP Instructs Skp2-dependent Cell Cycle Exit and Oncogenic Signaling .” The EMBO Journal 36 (17): 2510–28. https://doi.org/10.15252/embj.201696089.

47. Joshi, Jyotsna, Gautam Mahajan, and Chandrasekhar R. Kothapalli. 2018. “Three-Dimensional Collagenous Niche and Azacytidine Selectively Promote Time-Dependent Cardiomyogenesis from Human Bone Marrow-Derived MSC Spheroids.” Biotechnology and Bioengineering 115 (8): 2013–26. https://doi.org/10.1002/bit.26714.

48. Julian, Linda, and Michael F Olson. 2014. “Rho-Associated Coiled-Coil Containing Kinases (ROCK).” Small GTPases 5 (2): e29846. https://doi.org/10.4161/sgtp.29846.

49. Kamentsky, Lee, Thouis R Jones, Adam Fraser, Mark-Anthony Bray, David J Logan, Katherine L Madden, Vebjorn Ljosa, Curtis Rueden, Kevin W Eliceiri, and Anne E Carpenter. 2011. “Improved Structure, Function and Compatibility for CellProfiler: Modular High- Throughput Image Analysis Software.” *Bioinformatics (Oxford*, England) 27 (8): 1179–80. https://doi.org/10.1093/bioinformatics/btr095.

50. Kapoor, Avnish, Wantong Yao, Haoqiang Ying, Sujun Hua, Alison Liewen, Qiuyun Wang, Yi Zhong, et al. 2014. “Yap1 Activation Enables Bypass of Oncogenic KRAS Addiction in Pancreatic Cancer.” Cell 158 (1): 185–97. https://doi.org/10.1016/j.cell.2014.06.003.

51. Katoh, K, Y Kano, M Masuda, H Onishi, and K Fujiwara. 1998. “Isolation and Contraction of the Stress Fiber.” Molecular Biology of the Cell 9 (7): 1919–38. https://doi.org/10.1091/mbc.9.7.1919.

52. Kedersha, Nancy, and Paul Anderson. 2007. “Mammalian Stress Granules and Processing Bodies.” Methods in Enzymology 431 (07): 61–81. https://doi.org/10.1016/S0076-6879(07)31005-7.

53. Kedersha, Nancy, Sarah Tisdale, Tyler Hickman, and Paul Anderson. 2008. Chapter 26 Real- Time and Quantitative Imaging of Mammalian Stress Granules and Processing Bodies. Methods in Enzymology. 1st ed. Vol. 448. Elsevier Inc. https://doi.org/10.1016/S0076-6879(08)02626-8.

54. Kimura, K, M Ito, M Amano, K Chihara, Y Fukata, M Nakafuku, B Yamamori, et al. 1996. “Regulation of Myosin Phosphatase by Rho and Rho-Associated Kinase (Rho-Kinase).” *Science (New York*, N.Y.) 273 (5272): 245–48. https://doi.org/10.1126/science.273.5272.245.

55. Kimura, Masahiro, Takahiro Horie, Osamu Baba, Yuya Ide, Shuhei Tsuji, Randolph Ruiz Rodriguez, Toshimitsu Watanabe, et al. 2020. “Homeobox A4 Suppresses Vascular Remodeling by Repressing YAP/TEAD Transcriptional Activity.” EMBO Reports 21 (4): e48389. https://doi.org/10.15252/embr.201948389.

56. Kong, Fang, Andrés J García, A Paul Mould, Martin J Humphries, and Cheng Zhu. 2009. “Demonstration of Catch Bonds between an Integrin and Its Ligand.” Journal of Cell Biology 185 (7): 1275–84. https://doi.org/10.1083/jcb.200810002.

57. Kovacs, M, J Toth, C Hetenyi, A Malnasi-Csizmadia, and JR Sellers. 2004. “Mechanism of Blebbistatin Inhibition of Myosin II.” Journal of Biological Chemistry 279 (34): 35557–63. https://doi.org/10.1074/jbc.M405319200.

58. Lagunoff, M. 2015. KSHV-TK: Thymidine kinase or tyrosine kinase? EMBO J. 34 (4): 427–9.

59. Lai, Jason Kuan Han, and Didier Y.R. Stainier. 2017. “Pushing Yap into the Nucleus with Shear Force.” Developmental Cell 40 (6): 517–18. https://doi.org/10.1016/j.devcel.2017.03.008.

60. Lazarides, Elias, and Keith Burridge. 1975. “α-Actinin: Immunofluorescent Localization of a Muscle Structural Protein in Nonmuscle Cells.” Cell 6 (3): 289–98. https://doi.org/10.1016/0092-8674(75)90180-4.

61. Lee, Byoungkoo, Xin Zhou, Kristin Riching, Kevin W Eliceiri, Patricia J Keely, Scott A Guelcher, Alissa M Weaver, and Yi Jiang. 2014. “A Three-Dimensional Computational Model of Collagen Network Mechanics.” Edited by Sanjay Kumar. PLoS ONE 9 (11): e111896. https://doi.org/10.1371/journal.pone.0111896.

62. Lee, Hyun Jung, Miguel F Diaz, Katherine M Price, Joyce A Ozuna, Songlin Zhang, Eva M Sevick-Muraca, John P. Hagan, and Pamela L. Wenzel. 2017. “Fluid Shear Stress Activates YAP1 to Promote Cancer Cell Motility.” Nature Communications 8 (1): 14122. https://doi.org/10.1038/ncomms14122.

63. Lee, JSH. 2006. “Ballistic Intracellular Nanorheology Reveals ROCK-Hard Cytoplasmic Stiffening Response to Fluid Flow.” Journal of Cell Science 119 (9): 1760–68. https://doi.org/10.1242/jcs.02899.

64. Lee, Stacey, and Sanjay Kumar. 2016. “Actomyosin Stress Fiber Mechanosensing in 2D and 3D.” F1000Research 5 (0): 2261. https://doi.org/10.12688/f1000research.8800.1.

65. Li, Ying, Rui Chen, Qian Zhou, Zhisheng Xu, Chao Li, Shuai Wang, Aiping Mao, Xiaodong Zhang, Weiwu He, and Hong Bing Shu. 2012. “LSm14A Is a Processing Body-Associated Sensor of Viral Nucleic Acids That Initiates Cellular Antiviral Response in the Early Phase of Viral Infection.” Proceedings of the National Academy of Sciences of the United States of America 109 (29): 11770–75. https://doi.org/10.1073/pnas.1203405109.

66. Liu, G, FX Yu, YC Kim, Z. Meng, J Naipauer, DJ Looney, X Liu, JS Gutkind, EA Mesri, and K L Guan. 2015. “Kaposi Sarcoma-Associated Herpesvirus Promotes Tumorigenesis by Modulating the Hippo Pathway.” Oncogene 34 (27): 3536–46. https://doi.org/10.1038/onc.2014.281.

67. Liu, Huan, Xiaoming Dai, Xiaolei Cao, Huan Yan, Xinyan Ji, Haitao Zhang, Shuying Shen, et al. 2018. “PRDM4 Mediates YAP-induced Cell Invasion by Activating Leukocyte-specific Integrin Β2 Expression.” EMBO Reports 19 (6): 1–14. https://doi.org/10.15252/embr.201745180.

68. Lumb, Jennifer H, Lauren M Popov, Siyuan Ding, Marie T Keith, Bryan D Merrill, Harry B Greenberg, Jan E. Carette, Qin Li, and Jin Billy Li. 2017. “DDX6 Represses Aberrant Activation of Interferon-Stimulated Genes.” Cell Reports 20 (4): 819–31. https://doi.org/10.1016/j.celrep.2017.06.085.

69. McCormick, Craig, and Don Ganem. 2005. “The Kaposin B Protein of KSHV Activates the P38/MK2 Pathway and Stabilizes Cytokine MRNAs.” Science 307 (5710): 739–41. https://doi.org/10.1126/science.1105779.

70. Mo, Jung Soon, Zhipeng Meng, Young Chul Kim, Hyun Woo Park, Carsten Gram Hansen, Soohyun Kim, Dae Sik Lim, and Kun Liang Guan. 2015. “Cellular Energy Stress Induces AMPK-Mediated Regulation of YAP and the Hippo Pathway.” Nature Cell Biology 17 (4): 500–510. https://doi.org/10.1038/ncb3111.

71. Montaner, Silvia, Akrit Sodhi, Amanda K. Ramsdell, Daniel Martin, Jiadi Hu, Earl T Sawai, and J. Silvio Gutkind. 2006. “The Kaposi’s Sarcoma-Associated Herpesvirus G Protein-Coupled Receptor as a Therapeutic Target for the Treatment of Kaposi’s Sarcoma.” Cancer Research 66 (1): 168–74. https://doi.org/10.1158/0008-5472.CAN-05-1026.

72. Moreno-Vicente, Roberto, Dácil María Pavón, Inés Martín-Padura, Mauro Català-Montoro, Alberto Díez-Sánchez, Antonio Quílez-Álvarez, Juan Antonio López, et al. 2018. “Caveolin-1 Modulates Mechanotransduction Responses to Substrate Stiffness through Actin-Dependent Control of YAP.” Cell Reports 25 (6): 1622–1635.e6. https://doi.org/10.1016/j.celrep.2018.10.024.

73. Nakajima, Hiroyuki, and Naoki Mochizuki. 2017. “Flow Pattern-Dependent Endothelial Cell Responses through Transcriptional Regulation.” Cell Cycle 16 (20): 1893–1901. https://doi.org/10.1080/15384101.2017.1364324.

74. Nakajima, Hiroyuki, Kimiko Yamamoto, Sobhika Agarwala, Kenta Terai, Hajime Fukui, Shigetomo Fukuhara, Koji Ando, et al. 2017. “Flow-Dependent Endothelial YAP Regulation Contributes to Vessel Maintenance.” Developmental Cell 40 (6): 523–536.e6. https://doi.org/10.1016/j.devcel.2017.02.019.

75. Naranatt, Pramod P, Shaw M. Akula, Christopher A Zien, Harinivas H Krishnan, and Bala Chandran. 2003. “Kaposi’s Sarcoma-Associated Herpesvirus Induces the Phosphatidylinositol 3-Kinase-PKC-ζ-MEK-ERK Signaling Pathway in Target Cells Early during Infection: Implications for Infectivity.” Journal of Virology 77 (2): 1524–39. https://doi.org/10.1128/jvi.77.2.1524-1539.2003.

76. Ng, Chen Seng, Dacquin M Kasumba, Takashi Fujita, and Honglin Luo. 2020. “Spatio-Temporal Characterization of the Antiviral Activity of the XRN1-DCP1/2 Aggregation against Cytoplasmic RNA Viruses to Prevent Cell Death.” Cell Death and Differentiation. https://doi.org/10.1038/s41418-020-0509-0.

77. Noria, Sabrena, Feng Xu, Shannon McCue, Mara Jones, Avrum I Gotlieb, and B Lowell Langille. 2004. “Assembly and Reorientation of Stress Fibers Drives Morphological Changes to Endothelial Cells Exposed to Shear Stress.” The American Journal of Pathology 164 (4): 1211–23. https://doi.org/10.1016/S0002-9440(10)63209-9.

78. Núñez, Rocío Daviña, Matthias Budt, Sandra Saenger, Katharina Paki, Ulrike Arnold, Anne Sadewasser, and Thorsten Wolff. 2018. “The RNA Helicase DDX6 Associates with RIG-I to Augment Induction of Antiviral Signaling.” International Journal of Molecular Sciences 19 (7): 1–14. https://doi.org/10.3390/ijms19071877.

79. Ohashi, Kazumasa, Kyoko Nagata, Midori Maekawa, Toshimasa Ishizaki, Shuh Narumiya, and Kensaku Mizuno. 2000. “Rho-Associated Kinase ROCK Activates LIM-Kinase 1 by Phosphorylation at Threonine 508 within the Activation Loop.” Journal of Biological Chemistry 275 (5): 3577–82. https://doi.org/10.1074/jbc.275.5.3577.

80. Ojala, Päivi M, and Thomas F Schulz. 2014. “Manipulation of Endothelial Cells by KSHV: Implications for Angiogenesis and Aberrant Vascular Differentiation.” Seminars in Cancer Biology 26 (June): 69–77. https://doi.org/10.1016/j.semcancer.2014.01.008.

81. Ostareck, Dirk H, Isabel S Naarmann-de Vries, and Antje Ostareck-Lederer. 2014. “DDX6 and Its Orthologs as Modulators of Cellular and Viral RNA Expression.” Wiley Interdisciplinary Reviews: RNA 5 (5): 659–78. https://doi.org/10.1002/wrna.1237.

82. Panciera, Tito, Luca Azzolin, Atsushi Fujimura, Daniele Di Biagio, Chiara Frasson, Silvia Bresolin, Sandra Soligo, et al. 2016. “Induction of Expandable Tissue-Specific Stem/Progenitor Cells through Transient Expression of YAP/TAZ.” Cell Stem Cell 19 (6): 725–37. https://doi.org/10.1016/j.stem.2016.08.009.

83. Pavel, Mariana, Maurizio Renna, So Jung Park, Fiona M Menzies, Thomas Ricketts, Jens Füllgrabe, Avraham Ashkenazi, et al. 2018. “Contact Inhibition Controls Cell Survival and Proliferation via YAP/TAZ-Autophagy Axis.” Nature Communications 9 (1). https://doi.org/10.1038/s41467-018-05388-x.

84. Pellegrin, S, and H Mellor. 2007. “Actin Stress Fibres.” Journal of Cell Science 120 (20): 3491– 99. https://doi.org/10.1242/jcs.018473.

85. Pollard, Thomas D. 2016. “Actin and Actin-Binding Proteins.” Cold Spring Harbor Perspectives in Biology 8 (8): a018226. https://doi.org/10.1101/cshperspect.a018226.

86. Ridley, Anne J, and Alan Hall. 1992. “The Small GTP-Binding Protein Rho Regulates the Assembly of Focal Adhesions and Actin Stress Fibers in Response to Growth Factors.” Cell 70 (3): 389–99. https://doi.org/10.1016/0092-8674(92)90163-7.

87. Rio, Armando del, Raul Perez-Jimenez, Ruchuan Liu, Pere Roca-Cusachs, Julio M Fernandez, and Michael P Sheetz. 2009. “Stretching Single Talin Rod.” Science 323 (5914): 638–41.

88. Russo, James J, Roy A Bohenzky, MC Chien, Jing Chen, Ming Yan, Dawn Maddalena, J Preston Parry, et al. 1996. “Nucleotide Sequence of the Kaposi Sarcoma-Associated Herpesvirus (HHV8).” Proceedings of the National Academy of Sciences 93 (25): 14862–67. https://doi.org/10.1073/pnas.93.25.14862.

89. Schmitz, Arndt AP, Eve-Ellen Govek, Benjamin Böttner, and Linda Van Aelst. 2000. “Rho GTPases: Signaling, Migration, and Invasion.” Experimental Cell Research 261 (1): 1–12. https://doi.org/10.1006/excr.2000.5049.

90. Sharma, Nishi R, Vladimir Majerciak, Michael J Kruhlak, Lulu Yu, Jeong Gu Kang, Acong Yang, Shuo Gu, Marvin J Fritzler, and Zhi-ming Zheng. 2019. “KSHV RNA-Binding Protein ORF57 Inhibits P-Body Formation to Promote Viral Multiplication by Interaction with Ago2 and GW182,” 1–18. https://doi.org/10.1093/nar/gkz683.

91. Shaw, Gray, and Robert Kamen. 1986. “A Conserved AU Sequence from the 3′ Untranslated Region of GM-CSF MRNA Mediates Selective MRNA Degradation.” Cell 46 (5): 659–67. https://doi.org/10.1016/0092-8674(86)90341-7.

92. Shen, Zhewei, and Ben Z Stanger. 2015. “YAP Regulates S-Phase Entry in Endothelial Cells.” Edited by Deanna M Koepp. PLOS ONE 10 (1): e0117522. https://doi.org/10.1371/journal.pone.0117522.

93. Shepard, LW, M. Yang, P Xie, DD Browning, T Voyno-Yasenetskaya, T Kozasa, RD Ye. 2001. Constitutive activation of NF-kappa B and secretion of interleukin-8 induced by the G protein-coupled receptor of Kaposi’s sarcoma-associated herpesvirus involve G alpha(13) and RhoA. J Biol Chem. 276 (49):45979–87.

94. Sheth, Ujwal. 2003. “Decapping and Decay of Messenger RNA Occur in Cytoplasmic Processing Bodies.” Science 300 (5620): 805–8. https://doi.org/10.1126/science.1082320.

95. Shyu, AB, ME Greenberg, and JG Belasco. 1989. “The C-Fos Transcript Is Targeted for Rapid Decay by Two Distinct MRNA Degradation Pathways.” Genes & Development 3 (1): 60– 72. https://doi.org/10.1101/gad.3.1.60.

96. Small, J Victor, K Rottner, I Kaverina, and KI Anderson. 1998. “Assembling an Actin Cytoskeleton for Cell Attachment and Movement.” Biochimica et Biophysica Acta - Molecular Cell Research 1404 (3): 271–81. https://doi.org/10.1016/S0167-4889(98)00080-9.

97. Speck, Samuel H, and Don Ganem. 2010. “Viral Latency and Its Regulation: Lessons from the γ-Herpesviruses.” Cell Host and Microbe 8 (1): 100–115. https://doi.org/10.1016/j.chom.2010.06.014.

98. Staskus, K A, W Zhong, K Gebhard, B Herndier, H Wang, R Renne, J Beneke, et al. 1997. “Kaposi’s Sarcoma-Associated Herpesvirus Gene Expression in Endothelial (Spindle) Tumor Cells.” Journal of Virology 71 (1): 715–19. https://doi.org/10.1128/jvi.71.1.715-719.1997.

99. Stoecklin, Georg, Thomas Mayo, and Paul Anderson. 2006. “ARE-MRNA Degradation Requires the 5’-3’ Decay Pathway.” EMBO Reports 7 (1): 72–77. https://doi.org/10.1038/sj.embor.7400572.

100. Sugimoto, Wataru, Katsuhiko Itoh, Yasumasa Mitsui, Takahiro Ebata, Hideaki Fujita, Hiroaki Hirata, and Keiko Kawauchi. 2018. “Substrate Rigidity-Dependent Positive Feedback Regulation between YAP and ROCK2.” Cell Adhesion and Migration 12 (2): 101–8. https://doi.org/10.1080/19336918.2017.1338233.

101. Takahashi, Shinya, Kyoko Sakurai, Arisa Ebihara, Hiroaki Kajiho, Kota Saito, Kenji Kontani, Hiroshi Nishina, and Toshiaki Katada. 2011. “RhoA Activation Participates in Rearrangement of Processing Bodies and Release of Nucleated AU-Rich MRNAs.” Nucleic Acids Research 39 (8): 3446–57. https://doi.org/10.1093/nar/gkq1302.

102. Tan, John L, Joe Tien, Dana M Pirone, Darren S Gray, Kiran Bhadriraju, and Christopher S Chen. 2003. “Cells Lying on a Bed of Microneedles: An Approach to Isolate Mechanical Force.” Proceedings of the National Academy of Sciences 100 (4): 1484–89. https://doi.org/10.1073/pnas.0235407100.

103. Tojkander, Sari, Gergana Gateva, and Pekka Lappalainen. 2012. “Actin Stress Fibers - Assembly, Dynamics and Biological Roles.” Journal of Cell Science 125 (Pt 8): 1855–64. https://doi.org/10.1242/jcs.098087.

104. Tutucci, Evelina, Maria Vera, Jeetayu Biswas, Jennifer Garcia, Roy Parker, and Robert H. Singer. 2018. “An Improved MS2 System for Accurate Reporting of the MRNA Life Cycle.” Nature Methods 15 (1): 81–89. https://doi.org/10.1038/nmeth.4502.

105. Umbach, Jennifer L, Hui Ling Yen, Leo L M Poon, and Bryan R Cullen. 2010. “Influenza a Virus Expresses High Levels of an Unusual Class of Small Viral Leader RNAs in Infected Cells.” MBio 1 (4): 1–8. https://doi.org/10.1128/mBio.00204-10.

106. Vallenius, Tea. 2013. “Actin Stress Fibre Subtypes in Mesenchymal-Migrating Cells.” Open Biology 3 (6): 130001. https://doi.org/10.1098/rsob.130001.

107. Vassilev, Alex, Kotaro J Kaneko, Hongjun Shu, Yingming Zhao, and Melvin L. DePamphilis. 2001. “TEAD/TEF Transcription Factors Utilize the Activation Domain of YAP65, a Src/Yes-Associated Protein Localized in the Cytoplasm.” Genes and Development 15 (10): 1229–41. https://doi.org/10.1101/gad.888601.

108. Vindry, Caroline, Aline Marnef, Helen Broomhead, Laure Twyffels, Sevim Ozgur, Georg Stoecklin, Miriam Llorian, et al. 2017. “Dual RNA Processing Roles of Pat1b via Cytoplasmic Lsm1-7 and Nuclear Lsm2-8 Complexes.” Cell Reports 20 (5): 1187–1200. https://doi.org/10.1016/j.celrep.2017.06.091.

109. Vozzi, Federico, Jonica Campolo, Lorena Cozzi, Gianfranco Politano, Stefano Di Carlo, Michela Rial, Claudio Domenici, and Oberdan Parodi. 2018. “Computing of Low Shear Stress-Driven Endothelial Gene Network Involved in Early Stages of Atherosclerotic Process.” BioMed Research International 2018 (September): 1–12. https://doi.org/10.1155/2018/5359830.

110. Wada, KI, K Itoga, T Okano, S Yonemura, and H Sasaki. 2011. “Hippo Pathway Regulation by Cell Morphology and Stress Fibers.” Development 138 (18): 3907–14. https://doi.org/10.1242/dev.070987.

111. Wang, Huanru, Liang Chang, Xiaohui Wang, Airong Su, Chunhong Feng, Yuxuan Fu, Deyan Chen, Nan Zheng, and Zhiwei Wu. 2016. “MOV10 Interacts with Enterovirus 71 Genomic 5′UTR and Modulates Viral Replication.” Biochemical and Biophysical Research Communications 479 (3): 571–77. https://doi.org/10.1016/j.bbrc.2016.09.112.

112. Wang, Kuei-Chun, Yi-Ting Yeh, Phu Nguyen, Elaine Limqueco, Jocelyn Lopez, Satenick Thorossian, Kun-Liang Guan, Yi-Shuan J. Li, and Shu Chien. 2016. “Flow-Dependent YAP/TAZ Activities Regulate Endothelial Phenotypes and Atherosclerosis.” Proceedings of the National Academy of Sciences 113 (41): 11525–30. https://doi.org/10.1073/pnas.1613121113.

113. Watanabe, Naoki, Takayuki Kato, Akiko Fujita, Toshimasa Ishizaki, and Shuh Narumiya. 1999. “Cooperation between MDia1 and ROCK in Rho-Induced Actin Reorganization.” Nature Cell Biology 1 (3): 136–43. https://doi.org/10.1038/11056.

114. Watanabe, Naoki, Pascal Madaule, Tim Reid, Toshimasa Ishizaki, Go Watanabe, Akira Kakizuka, Yuji Saito, Kazuwa Nakao, Brigitte M. Jockusch, and Shuh Narumiya. 1997. “P140mDia, a Mammalian Homolog of Drosophila Diaphanous, Is a Target Protein for Rho Small GTPase and Is a Ligand for Profilin.” EMBO Journal 16 (11): 3044–56. https://doi.org/10.1093/emboj/16.11.3044.

115. Wilbertz, Johannes H, Franka Voigt, Ivana Horvathova, Gregory Roth, Yinxiu Zhan, and Jeffrey A. Chao. 2019. “Single-Molecule Imaging of MRNA Localization and Regulation during the Integrated Stress Response.” Molecular Cell 73 (5): 946–958.e7. https://doi.org/10.1016/j.molcel.2018.12.006.

116. Winzen, Reinhard, Michael Kracht, Birgit Ritter, Arno Wilhelm, Chyi Ying A Chen, Ann Bin Shyu, Monika Müller, Matthias Gaestel, Klaus Resch, and Helmut Holtmann. 1999. “The P38 MAP Kinase Pathway Signals for Cytokine-Induced MRNA Stabilization via MAP Kinase-Activated Protein Kinase 2 and an AU-Rich Region-Targeted Mechanism.” EMBO Journal 18 (18): 4969–80. https://doi.org/10.1093/emboj/18.18.4969.

117. Yang, Chih-Sheng, Eleni Stampouloglou, Nathan M Kingston, Liye Zhang, Stefano Monti, and Xaralabos Varelas. 2018. “Glutamine-utilizing Transaminases Are a Metabolic Vulnerability of TAZ/YAP-activated Cancer Cells.” EMBO Reports 19 (6): 1–11. https://doi.org/10.15252/embr.201643577.

118. Yang, Ya Li, Lindsay M. Leone, and Laura J. Kaufman. 2009. “Elastic Moduli of Collagen Gels Can Be Predicted from Two-Dimensional Confocal Microscopy.” Biophysical Journal 97 (7): 2051–60. https://doi.org/10.1016/j.bpj.2009.07.035.

119. Yu, Fa Xing, Bin Zhao, Nattapon Panupinthu, Jenna L. Jewell, Ian Lian, Lloyd H. Wang, Jiagang Zhao, et al. 2012. “Regulation of the Hippo-YAP Pathway by G-Protein-Coupled Receptor Signaling.” Cell 150 (4): 780–91. https://doi.org/10.1016/j.cell.2012.06.037.

120. Yu, Jiang Hong, Wei-Hong Yang, Tod Gulick, Kenneth D Bloch, and Donald B Bloch. 2005. “Ge-1 Is a Central Component of the Mammalian Cytoplasmic MRNA Processing Body.” *RNA (New York*, N.Y.) 11 (12): 1795–1802. https://doi.org/10.1261/rna.2142405.

121. Zanconato, Francesca, Mattia Forcato, Giusy Battilana, Luca Azzolin, Erika Quaranta, Beatrice Bodega, Antonio Rosato, Silvio Bicciato, Michelangelo Cordenonsi, and Stefano Piccolo. 2015. “Genome-Wide Association between YAP/TAZ/TEAD and AP-1 at Enhancers Drives Oncogenic Growth.” Nature Cell Biology 17 (9): 1218–27. https://doi.org/10.1038/ncb3216.

122. Zhang, Qian, Fansen Meng, Shasha Chen, Steven W Plouffe, Shiying Wu, Shengduo Liu, Xinran Li, et al. 2017. “Hippo Signalling Governs Cytosolic Nucleic Acid Sensing through YAP / TAZ-Mediated TBK1 Blockade.” Nature Cell Biology 19 (4): 362–74. https://doi.org/10.1038/ncb3496.

123. Zhao, Bin, Li Li, Lloyd Wang, Cun Yu Wang, Jindan Yu, and Kun Liang Guan. 2012. “Cell Detachment Activates the Hippo Pathway via Cytoskeleton Reorganization to Induce Anoikis.” Genes and Development 26 (1): 54–68. https://doi.org/10.1101/gad.173435.111.

124. Zhao, Bin, Bin Zhao, Xiaomu Wei, Xiaomu Wei, Weiquan Li, Weiquan Li, Ryan S Udan, et al. 2007. “Inactivation of YAP Oncoprotein by the Hippo Pathway Is Involved in Cell Contact Inhibition and Tissue Growth Control.” Genes and Development 21: 2747–61. https://doi.org/10.1101/gad.1602907.Hpo/Sav.

125. Zhong, W, H Wang, B Herndier, and D Ganem. 1996. “Restricted Expression of Kaposi Sarcoma-Associated Herpesvirus (Human Herpesvirus 8) Genes in Kaposi Sarcoma.” Proceedings of the National Academy of Sciences of the United States of America 93 (13): 6641–46. https://doi.org/10.1073/pnas.93.13.6641.

